# Advancing Glycan Analysis: A New Platform Integrating SERS, Boronic Acids, and Machine Learning Algorithms

**DOI:** 10.1101/2023.02.03.527030

**Authors:** Qiang Hu, Dacheng Kuai, Hyundo Park, Haley Clark, Perla B. Balbuena, Joseph Sang-Il Kwon, Hung-Jen Wu

## Abstract

Glycans are the most abundant and diverse fundamental biomolecules. Profiling glycans is essential to elucidate biomolecular mechanisms and to predict biological consequences; however, the analysis of glycans remains challenging due to their structural complexity. A novel glycan detection platform is established through the integration of surface-enhanced Raman spectroscopy (SERS), boronic acid receptors, and machine learning tools. SERS is able to distinguish isomeric structures and provide fingerprint information of molecules. Boronic acid receptors can selectively bind with glycans, and the reaction influences molecular vibrations, leading to unique Raman spectral patterns. For proof-of-concept, two boronic acids, 4-mercaptophenylboronic acid (4MBA) and 1-thianthrenylboronic acid (1TBA) were used for glycan detection. The spectra were analyzed by the machine learning algorithm. The sensing platform successfully recognized the stereoisomers (glucose, mannose, and galactose), the structural isomers with α-1,4 and α-1,6 glycosidic linkages, and the β-1,4 glycosidic linkage within lactose molecules. The collective spectra that combine the spectra from both boronic acid receptors could improve the performance of the support vector machine model due to the enrichment of the structural information of glycans. In addition to qualitative analysis, this new sensor could quantify the mole fraction of sialic acid in lactose background using the machine learning regression technique. The quantification accuracy reached a coefficient of determination (R^2^) value of 0.998 and a normalized mean square error (NMSE) of 0.00195. This low-cost, rapid, and highly accessible sensor will provide the scientific community with another option for frequent glycan screening in standard biological laboratories.

## Introduction

Glycans are the most abundant and diverse fundamental biomolecules in living organisms. Almost all cells in nature are covered with dense layers of glycans that mediate a wide range of biochemical reactions^1^. Because glycosylation is a post-translational modification (PTM), glycan profiles could be influenced by various factors, such as the environment, temperature, cell growth cycles, cell health, etc. As such, frequent monitoring of the glycosylation changes is needed to explore cell activities. Currently, the analysis of glycans remains challenging due to the complex nature seen from their isomeric glycan forms, glycosidic linkages, and branched structures. The common glycan analysis techniques, including mass spectrometry (MS) and nuclear magnetic resonance (NMR), are resource-intensive and typically require experienced scientists for operation and data analysis, making these techniques unsuitable for frequent screening in standard biological laboratories^2–4^. Due to these challenges, an accessible glycan analysis tool that allows users to frequently monitor the changes of the glycosylation is highly desirable^2, 4^.

Herein, we designed a new glycan sensing platform by integrating surface-enhanced Raman spectroscopy (SERS), boronic acid receptors, and machine learning techniques. Raman spectroscopy that offers fingerprint spectra of molecules is a potential tool for glycan analysis, but the inherent weak signals limit its applications. To overcome this barrier, a Raman enhancement technique, called SERS, was used to increase the Raman signals of molecules that are adsorbed on metallic nanoparticles^5^. In this study, a SERS substrate, named nanopaper, which is a glass fiber paper decorated with a dense layer of silver nanoparticles, was used for glycan detection^6–9^.

In addition, the SERS substrate was further functionalized with boronic acid to improve detection accuracy. Boronic acids have been widely used for the separation and immobilization of glycans because of the covalent reaction between boronic acids and cis-diols on glycans molecules^10^. Boronic acid coated nanoparticles have been used to capture glycan molecules^11–13^. By designing intermolecular distances between boronic acid moieties, boronic acids could selectively bind to different types of glycans^14^. Researchers have used different types of boronic acids to detect sialic acids in cancer cells^11, 15^, glycoproteins in serum samples^16, 17^, and glycans on cell surfaces^18, 19^. In addition, boronic acid analogs are typically attached to aryl groups that can offer fluorescent signals and distinct Raman vibrations^20^. We hypothesized that the reaction between cis-diols and different boronic acid analogs would cause different types of spectral shifts. By integrating the spectral information collected from different boronic acids, the accuracy of glycan identification may be improved.

Although Raman spectroscopy could offer the fingerprint spectra of glycan samples, the data interpretation remains challenging. Each glycan-boronic acid reaction pair would offer a few significant Raman peaks. The Raman peaks are likely to overlap when the glycan complexity increases. Machine learning-based methods have been used to solve complicated qualitative and quantitative questions^21, 22^. Same methods can be applied on Raman spectra since it could differentiate the spectra based on subtle changes and reveal the underlying data patterns from the whole spectra instead of individual peaks^23^. We integrated the discriminant analysis of principal components (DAPC) with various machine learning classifiers, including support vector machine (SVM), to classify SERS spectra collected from beverages^9^. The same machine learning strategy was applied to analyze the complex SERS spectra from various glycan samples.

For proof-of-concept, we studied the interactions of two boronic acids: 4-mercaptophenylboronic acid (4MBA) and 1-thianthrenylboronic acid (1TBA) with a group of monosaccharides and disaccharides (The molecular structures are shown in **Fig. S1**). The reaction between boronic acids and glycans could trigger distinctive spectral changes, and the experimental spectra agreed with the simulation data calculated by the density functional theory (DFT). The glycan-boronic acid reactions influenced the charge distributions of the aromatic structures on the boronic acids; thus, 1TBA and 4MBA receptors offered different types of glycan binding spectra. By combining the SERS spectra from 1TBA and 4MBA receptors, the “collective spectra” could enrich the available structural information. The results show that such collective spectra could improve classification accuracy. To evaluate the limitation, the sensor was used to differentiate stereoisomers (glucose, mannose, and galactose), structure isomers with different types of glycosidic linkages (maltose vs. isomaltose), and the presence of glycosidic linkages (lactose vs. glucose and galactose mixture). The classification accuracy of the selected glycans could reach 99.6%. In addition to qualitative analysis, the quantification of mole fraction of sialic acids in lactose background was conducted using the machine learning regression technique. The quantification accuracy reached a coefficient of determination (R^2^) value of 0.998 and a normalized mean square error (NMSE) of 0.00195.

## Methods

### Materials

Glucose (Glu), galactose (Gal), mannose (Man), fucose (Fuc), N-acetylglucosamine (GlcNAc), N-acetylgalactosamine (GalNAc), N-acetylneuraminic acid (Neu5Ac or sialic acid), maltose, isomaltose, lactose, ammonia hydroxide solution (28%-30%), 4-mercaptophenylboronic acid (4MBA), and 1-thianthrenylboronic acid (1TBA) were purchased from Sigma Aldrich (Glycan and boronic acid structures are shown in **Fig. S1**). Glass microfiber papers (Grade: 934-AH) were acquired from Whatman. Silver nitrate (99.99995%) was purchased from Alfa Aesar. 2-propanol was purchased from Fisher Scientific. All chemicals were ACS grade or higher and used without further purification.

### Nanopaper fabrication

Nanopapers were fabricated as previously reported^6^. In brief, Tollen’s reagent containing 300 mM ammonia and 50 mM silver nitrate was prepared in a 2L glass beaker in a 55°C water bath. Glass microfiber papers were immersed in the solution, and 500 mM glucose solution was added to initiate the silver mirror reaction. After the reaction was complete, the filter papers were rinsed thoroughly with deionized water and 2-propanol. The resulting products, i.e., the nanopapers, were stored in 2-propanol. The storage container was covered with aluminum foil and placed in drawers to prevent light exposure.

### Substrate surface modification and glycan detection

Before surface modification, nanopapers were cut into a 1 cm × 0.5 cm rectangular shape. Nanopapers were coated with boronic acids by immersing substrates in 50 mM 1TBA or 0.1 mM 4MBA in methanol for 1 hour. For glycan detection, the boronic acid coated nanopaper was immersed in the aqueous solution containing 1 mM of the desired glycans for 1 hour. Before Raman measurement, the paper was dried in an oven at 75°C for 5 minutes.

### Raman measurements

Raman spectra were collected with a ThermoFisher Scientific DXR3 Raman microscope using laser excitation with a wavelength of 785 nm and an output power of 1 mW. This instrument was equipped with an Olympus BX41 optical microscope and a thermoelectrically cooled charge-coupled detector (ThermoFisher front-illuminated CCD system) with 1024 × 256 pixel format, operating at −70 °C. The signal was calibrated by an internal polystyrene standard and a 10× objective. The spot size was about 3.8 μm. 200 SERS spectra were collected with an exposure time of 1 s for 5 accumulations at different spots for each sample.

### Data processing of Raman spectra

The data analysis was performed using the same methodology reported in the previous study^9^. The data analysis scheme is shown in **Fig S2**. Briefly, the spectra were first processed using asymmetric least square (ALS) baseline correction. Then, baselined spectra were vector normalized and Savitzky–Golay smoothed (4^th^ order polynomial, with a frame size of 37). Finally, multivariate analysis techniques and classification algorithms were performed in the spectral range 400-1650 cm^−1^. Data processing was conducted with Matlab 2021b.

### Multivariate analysis and machine learning classification

Before applying classifiers, the smoothed spectra were processed using multivariate statistical analysis to reduce the complexity and extract the significant spectral features that explain the most variance. Discriminant analysis of principal components (DAPC) was used in this study^24^. Principal component analysis (PCA) was first applied to reduce the data complexity, and then, a supervised multivariate analysis, discriminant analysis, was used to further discriminate the dataset by correlating the variation in the data with the sample information.

After feature extraction, common machine learning classifiers were used to classify SERS spectra. Support vector machine (SVM) was selected because of its superior performance in Raman spectral analysis^9^. A 5-fold cross-validation was performed to assess the suitability of the classification algorithm^25^. In brief, the training and the validation sets were established by randomly selecting from the Raman spectra data. The training dataset was used to generate a classification model, and the model predicted the validation dataset to evaluate the performance. The cross-validation approach was repeated five times, wherein the validation set is consisting of 280 randomly selected SERS spectra in repetition for each monosaccharide case, 120 for each disaccharide structural isomers case, 80 for each different glycosidic linkage case, 440 for all sample and collective spectra cases. The performance of the model was evaluated by classification accuracies, sensitivity, and selectivity. The classification accuracy, sensitivity, and selectivity are defined as^26^:

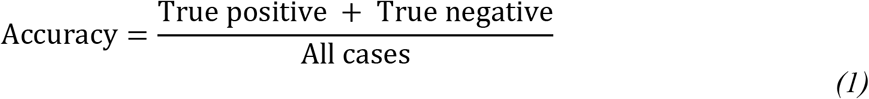

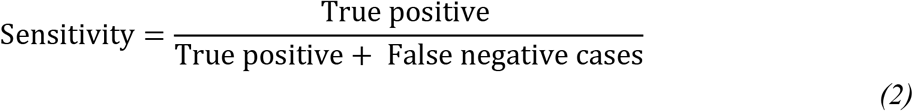

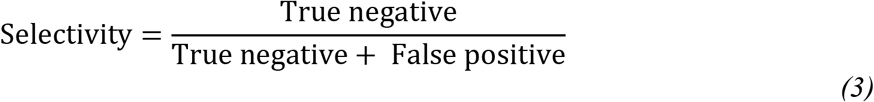

The collective spectra were constructed by combining the truncated 1TBA spectra (400-1650 cm^− 1^) and the 4MBA (400-1650 cm^−1^) spectra. Then, the collective spectra went through the same multivariate analysis and classification algorithm as the individual spectra.

### Sialic acid quantification

Different amounts of sialic acid were added to the 10 mM lactose solution to create a serial dilution. The final mole fractions of the sialic acid in the mixture are ranged from 0 mol.% to 2mol.%, with 0.5 mol.% intervals. The oligosaccharide mixtures were analyzed by SERS using the same protocol reported above. 200 spectra were collected for each concentration.

### Machine learning regression analysis

Regression analysis was conducted using Matlab 2021b. Gaussian process regression algorithm was used for predicting the mole fraction of sialic acid in lactose background, and the scheme is shown in **Fig. S2**. DAPC was first performed on the dataset for the wavenumber from 400-1650 cm^−1^, and then the resulting canonicals were used in regression analysis. A 5-fold cross-validation was performed on the model to evaluate the regression performance. The model was evaluated based on the normalized mean square error (NMSE) and coefficient of determination (R^2^). For the collective spectra regression, the dataset was built in the same way as described in the classification. Then the spectra went through the same regression algorithm and were evaluated based on the same performance metrics. A 5-fold cross-validation was performed on the model to evaluate the regression performance. The model was evaluated based on the normalized mean square error (NMSE) and coefficient of determination (R^2^).

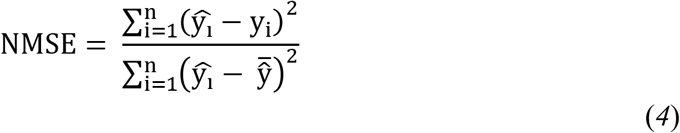

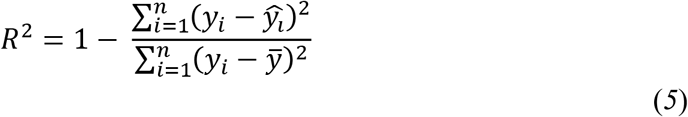

where *ŷ_i_* is the predicted values, *y_i_* is the actual values in the dataset, 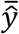 is the mean of the predicted values, *ȳ* is the mean of the actual values, and the n is the number of spectra in the dataset.

### DFT theoretical calculations

Structures of 4MBA, 1TBA, 1TBA with glucose-bound, and 1TBA with mannose-bound molecules were optimized at B3LYP level of theory with dipole-polar-enhanced 6-311++G(d,p) basis sets embedded in Gaussian 16 package^27^. Empirical dispersion correction was applied for better descriptions of hydrogen bonding^28^. Calculated Raman wavenumbers were calibrated based on the experimental data retrieved from NIST^29^. The vibration modes were assigned using the literature values, and unknowns were estimated using VEDA^12, 13, 30–32^.

## Results and Discussion

### Boronic acid coated SERS substrate

Glass fiber filter papers coated with a dense layer of silver nanoparticles were used as SERS substrates. This nanoplasmonic substrate, called nanopaper, was prepared by the silver mirror reaction, offering a significant Raman signal enhancement^6^. The SERS spectra of glucose, galactose, mannose, and sialic acid on the nanopaper without boronic acid surface modification are shown in **Fig. S3**. Glucose, galactose, and mannose are stereoisomers; the spectral difference among these isomers could be clearly observed. However, the intensities of Raman spectra are still relatively low on the nanopaper. SERS is a near-field phenomenon that enhances Raman signals near metal nanoparticle surfaces^33^. Less than desirable contacts between sugars and silver particles probably led to lower signals. Decorating additional receptor molecules on nanopapers could bring glycans near silver particle surfaces, eventually improving SERS signals^34^. The reporter molecule chosen here is boronic acids due to their covalent reactions towards cis-diols on glycans molecules. Boronic acids could selectively bind to various pairs of hydroxyl groups on glycans, based on intermolecular distances of boronic acid moieties. Thus, the nanopaper surfaces were modified with boronic acid molecules to capture glycan molecules.

The sensor design is shown in **Fig. 1**. We modified nanopaper surface with two types of boronic acids, 4MBA and 1TBA, and evaluated their performance. 4MBA has been extensively used for glycan sensor developments^11–13, 16, 31^, but 1TBA has not been thoroughly investigated. Compared to 4MBA, 1TBA has two aromatic rings in its structure. The spectral shifts caused by 1TBA-glycan binding would differ from those caused by 4MBA-glycan. We hypothesized that the collection of spectral information from different boronic acids can improve the accuracy of glycan identification.

**Figure 1.**
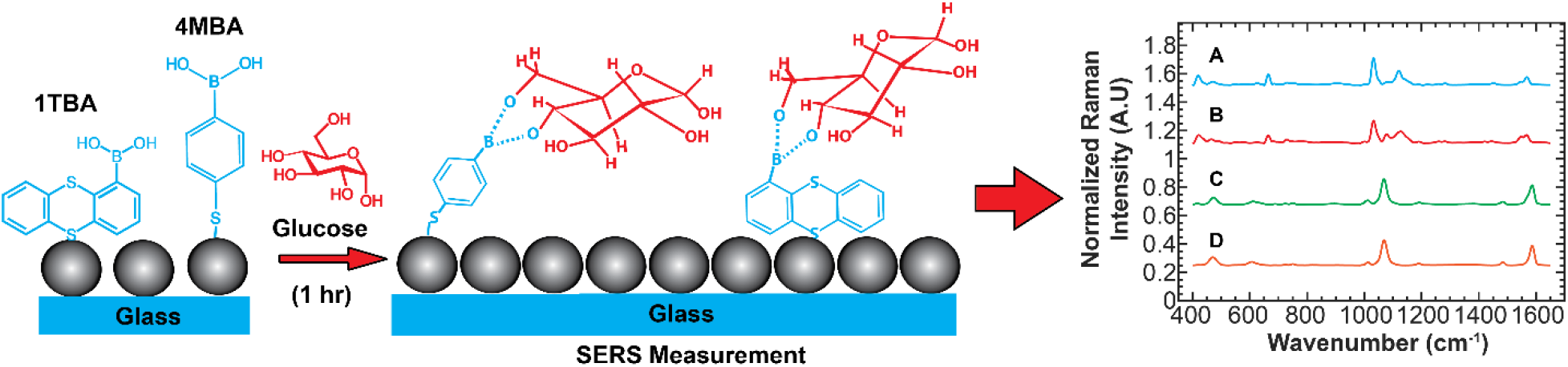
A schematic of the sensor design. SERS was conducted on nanopaper substrates (i.e., glass fiber papers decorated by silver nanoparticles). The surfaces of silver particles were modified by boronic acid receptors that could bind to hydroxyl groups on glycan molecules. The interactions between boronic acids and glycans influence molecular vibrations, leading to unique Raman spectral patterns. The complex Raman data were further analyzed by the machine learning algorithms for glycan classification and quantification. The example spectra shown here are (A) 1TBA with glucose, (B) 1TBA, (C) 4MBA with glucose, (D) 4MBA.

### Comparison of experimental and theoretical spectra

Raman scattering of 4MBA interacting with glycans has been studied extensively^12, 13, 31, 35^, and the vibrational assignments from the literature are listed in **Table S1**. 1TBA has not been fully investigated; thus, we measured and calculated the Raman spectra of 1TBA-glycan complex. **Fig. 2** shows the experimental SERS spectra of glucose and mannose on 1TBA functionalized nanopaper. The SERS signals are over 20 times stronger than the signals on the nanopaper without boronic acid functionalization. The spectral difference among no sugar (1TBA only), 1TBA-glucose, 1TBA-galactose, and 1TBA-mannose could be observed in **Fig. 2**.

**Figure 2.**
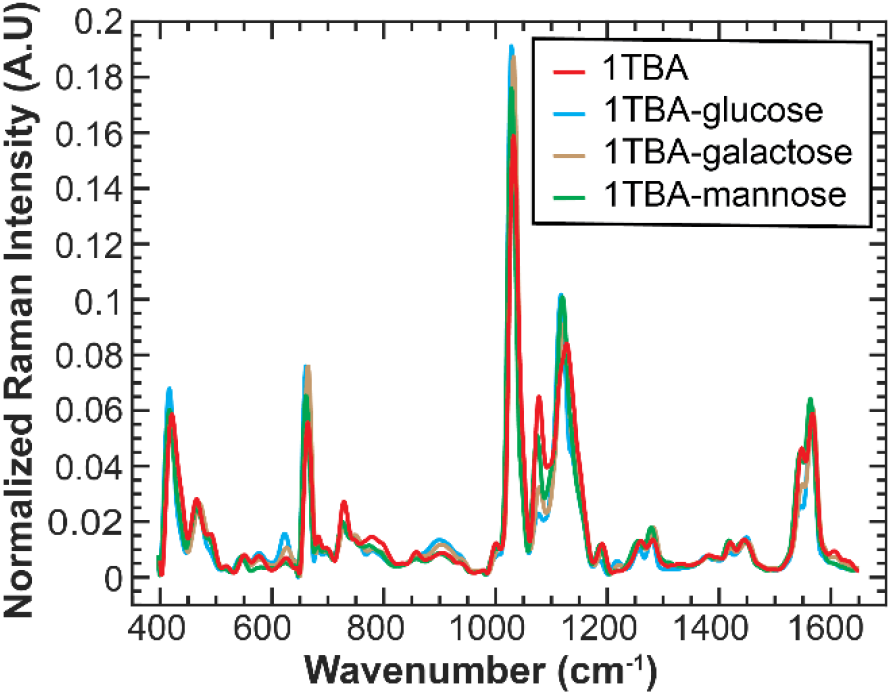
SERS spectra of 1TBA interacting with selected monosaccharides. The 1TBA spectra (control, red), 1TBA-glucose spectra (blue), 1TBA-galactose (brown) and 1TBA-mannose spectra (green) are shown here. Spectral differences among these three stereoisomers were observed.

Density functional theory (DFT) simulations were performed to understand the influence of molecular vibrations on Raman spectra. We conducted the simulations for 1TBA along and 1TBA binding to two stereoisomers, glucose and mannose. The prior study has shown that boronic acid derivatives could bind to various combinations of hydroxyl groups on monosaccharides^36^. The flexible hydroxyls on positions 4 and 6 of glucose are most likely to interact with boronic acids^37, 38^; thus, we conducted the DFT calculations of 1TBA binding to 4, 6 diols on glucose and mannose. The experimental and theoretical vibrational frequencies, along with corresponding vibrational assignments and intensities are shown in **Table S2**. The DFT simulation suggests that the interactions between 1TBA and monosaccharides influence the charge distributions in 1TBA aromatic ring structure (**Fig. S4**). In addition, the stereoisomer, glucose and mannose, could cause different charge distributions, leading to changes in Raman spectra. These findings correspond to the frontier molecular orbital (FMO) analysis done by Revanna et al^30^. For instance, DFT calculations show that the bindings of glucose and mannose with 1TBA could intensify the peak at 428 cm^−1^ (CCCC torsion and SCCC out of plane bending); moreover, 1TBA-glucose has a higher intensity at 421 cm^−1^ compared to 1TBA-mannose at 423 cm^−1^. (**Movie 1 in the supporting information**). The changes of simulated molecular vibrations can be observed in other Raman peaks as well, such as 1560 cm^−1^ and 1568 cm^−1^ (**Movie 2 & 3 in the supporting information)**. The changes in these theoretical values correspond to what we observed in the experimental results. The theoretical calculations support our hypothesis that the binding of glycans to boronic acid could shift the vibration modes, leading to unique changes in Raman spectra.

### Explore the spectral variations among seven common monosaccharides

We first evaluated the capability of monosaccharide detection. The common monosaccharides in mammalian cells, glucose, mannose, galactose, fucose, GlcNAc, GalNAc, and sialic acid, were investigated^39^. **Fig. 3** shows the SERS spectra of the selected monosaccharides using 4MBA or 1TBA functionalized nanopapers. SERS intensities from 4MBA functionalized nanopapers were stronger than the signals from 1TBA functionalized nanopapers. The thiol-silver bonding enhances the adsorption of 4MBA on silver nanoparticles, resulting in higher signal intensities^40^. Higher SERS signals of glucose-4MBA nanopapers lead to a higher signal-to-noise ratio (S/N ≈ 417), compared to the spectra on glucose-1TBA nanopapers (S/N ≈ 148).

**Figure 3.**
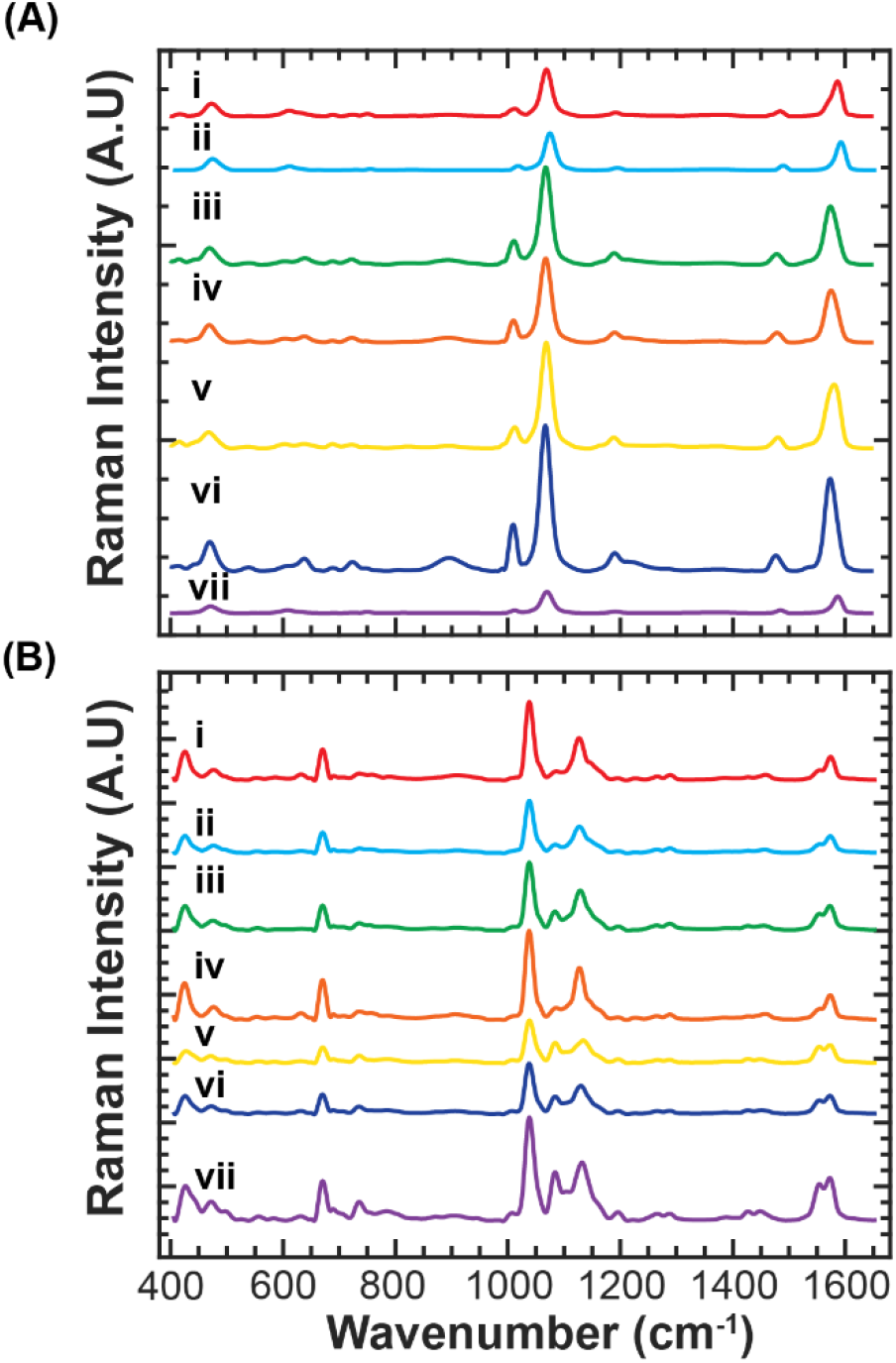
4MBA (A) and 1TBA (B) monosaccharide average SERS spectra. The spectra are glucose (i), galactose (ii), mannose (iii), GalNAc (iv), GlcNAc (v), fucose (vi), sialic acid (vii).

The principal component analysis (PCA) was used to explore the major spectral variations among the selected monosaccharides. The first PC explained 90% of the variabilities. The PCA contribution plots of PC1 for 4MBA and 1TBA nanopapers are shown in **Fig. S5**. For 4MBA nanopaper, one of the major changes in the vibrational spectra could be observed at 1008 cm^−1^, contributed by CC and OH stretching. The other major changes were located at 1570 cm^−1^ and 1589 cm^−1^, which are associated with CC stretching and CH bending on the aromatic ring of 4MBA. Compared with 4MBA spectra, more changes could be observed in 1TBA spectra, which likely means more vibration modes are influenced by the 1TBA-glycan binding process. For 1TBA, there is a shift from a 1095 cm^−1^ peak in 1TBA to an 1104 cm^−1^ peak in 1TBA-glucose and 1TBA-mannose SERS spectra (**Fig. 3**), which are CC stretching. The 1128 cm^−1^ peak in 1TBA has another shift to an 1139 cm^−1^ peak in 1TBA-glucose and 1TBA-mannose spectra, associated with CC stretching and HCC in-plane bending. These observations from 4MBA and 1TBA spectra show that the monosaccharide binding causes detectable changes in Raman spectra.

### Classification of monosaccharides via machine learning algorithm

PCA analysis indicated that the bindings of different monosaccharides to boronic acid functionalized nanopapers could cause spectral changes at various wavenumbers. The complex spectral variations could not be easily distinguished by visual inspection; thus, the multivariate analysis and machine learning algorithm were utilized to analyze the SERS spectra. SERS spectra were first processed by the multivariate analysis, DAPC, and then the classic machine learning classifiers, SVM, were used to classify the spectra. DAPC could extract spectral features related to the key difference among the sample groups and minimize the influence of the variations within the same sample groups^9, 24^. The combination of DAPC with SVM has shown superior performance for SERS spectra analysis^9^. The performance of the machine learning classification was expressed using the confusion matrix (**Fig. S6**).

The nanopapers functionalized with 4MBA show a better classification performance, compared to the nanopaper coated with 1TBA. For 4MBA nanopaper, the average classification accuracy could reach 99.7% and the average sensitivities and average specificities of the selected monosaccharide exceed 99.5%. For 1TBA nanopaper, the average classification accuracy is 97.4%, and the range of the sensitivities and specificities for selected monosaccharides are around 93~99%. 1TBA nanopaper particularly misclassified GlcNAc. The relatively poor performance of 1TBA nanopaper may be explained by a lower S/N. For the GlcNAc, the S/N on 4MBA and 1TBA nanopapers were 499 and 83.7, respectively. The acetyl group on the position C2 of GlcNAc may influence the interaction of GlcNAc with the borate groups, leading to a weaker binding affinity toward 1TBA. Nevertheless, both the boronic acids have the capability of distinguishing the selected monosaccharides, including the stereoisomers.

### Distinguish α-1,4 and α-1,6 glycosidic linkage in maltose and isomaltose

The types of glycosidic linkages between two saccharide units could influence biological function^41^. Identification of glycosidic linkages is essential in glycan detection. To evaluate the capability of the sensor, we first explore two disaccharides, maltose and isomaltose. Maltose and isomaltose are structural isomers composed of two glucose units. The glucose of maltose is connected via position 1 and 4, while isomaltose has a α-1,6 glycosidic linkage. **Fig. 4** shows the SERS spectra of maltose, isomaltose, and glucose using 4MBA or 1TBA functionalized nanopaper.

**Figure 4.**
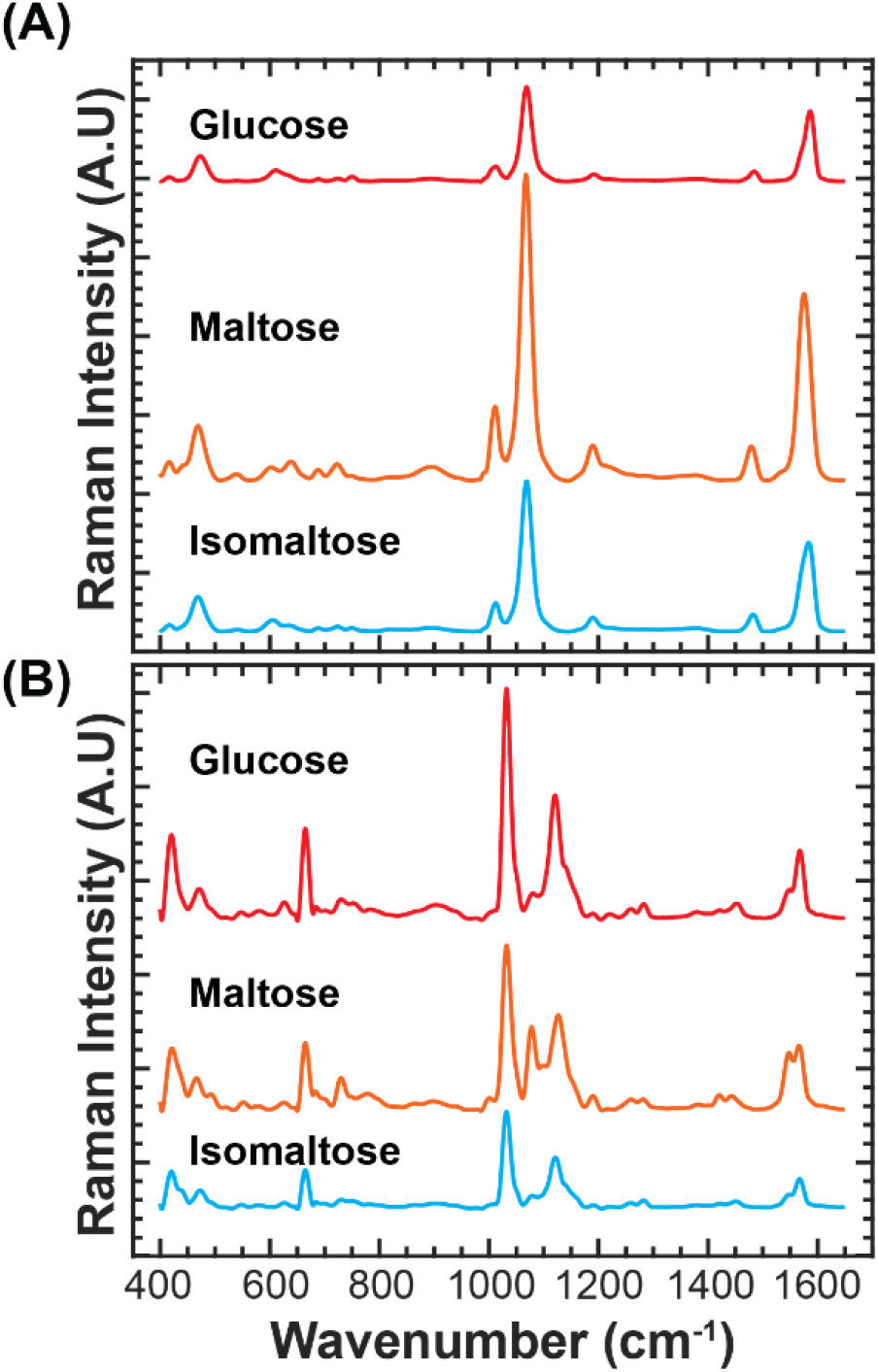
4MBA (A) and 1TBA (B) disaccharide average SERS spectra. The spectra are glucose, maltose, isomaltose.

PCA analysis was performed again to observe the major variations among those three samples. (**Fig. S7)**. The relative peak contributions from 4MBA are similar to the ones for monosaccharide cases. The 1570 cm^−1^ and 1589 cm^−1^ peaks contribute the most compared with other peaks, representing the CC stretching and CH bending on the aromatic ring structure of 4MBA. In the 1TBA spectra dataset **(Fig. S7. (B))**, the main contributions to distinguish the samples are from the 1078 cm^−1^, 1134 cm^−1^, 1547 cm^−1^, and 1564 cm^−1^ peaks, originating from the CC stretching and HCC in-plane bending. The 418 cm^−1^ peak for the SC stretching and SCC in-plane bending does not help as much in the disaccharide case compared to the monosaccharide case.

The machine learning classifier was used again to distinguish these three samples (**Fig. S8)**. The 4MBA and 1TBA functionalized nanopapers performed greatly in differentiating maltose, isomaltose, and glucose spectra with 99.8% and 100% accuracy, respectively.

### Distinguish glycosidic linkage between two stereoisomers, glucose and galactose

The capability of detecting another common disaccharide, lactose, was also evaluated in this study. Lactose consists of one glucose and one galactose with a β-1,4 linkage. Prior studies have suggested that the flexible hydroxyls on positions 4 and 6 of glucose are most likely to interact with boronic acids^37, 38^. This hydroxyl group of glucose at position 4 is blocked by the glycosidic linkage in lactose molecules. To observe the influence of the glycosidic bonds, we measured the SERS spectra of the lactose and an equal volume mixture of glucose and galactose. (**Fig. 5**) Upon visual inspection, the spectra of lactose and the mixture of glucose and galactose are very similar. PCA was performed to observe the tiny changes in SERS spectra. (**Fig. S9. (A)**) For 4MBA, the major spectral variations are similar to the changes in the prior monosaccharide, maltose, and isomaltose. The shifts of SERS peaks were observed at the 472 cm^−1^, 609 cm^−1^, and 1073 cm^−1^ regions. For 1TBA (**Fig. S9 (B)**), PC1 shows a different result compared with the previous 1TBA PC1 plots for monosaccharides and disaccharides. We found that PC1 has increased contributions from the 579 cm^−1^, 892 cm^−1^, 1341 cm^−1^, 1366 cm^−1^, 1417 cm^−1^, 1440 cm^−1^, 1452 cm^−1^, 1457 cm^−1^, 1468 cm^−1^, 1474 cm^−1^, 1487 cm^−1^, and 1493 cm^−1^ regions; however, these peaks exhibit lower signals. The low S/N of 1TBA receptor leads to the poor classification performance (**Fig. S10**).

**Figure 5.**
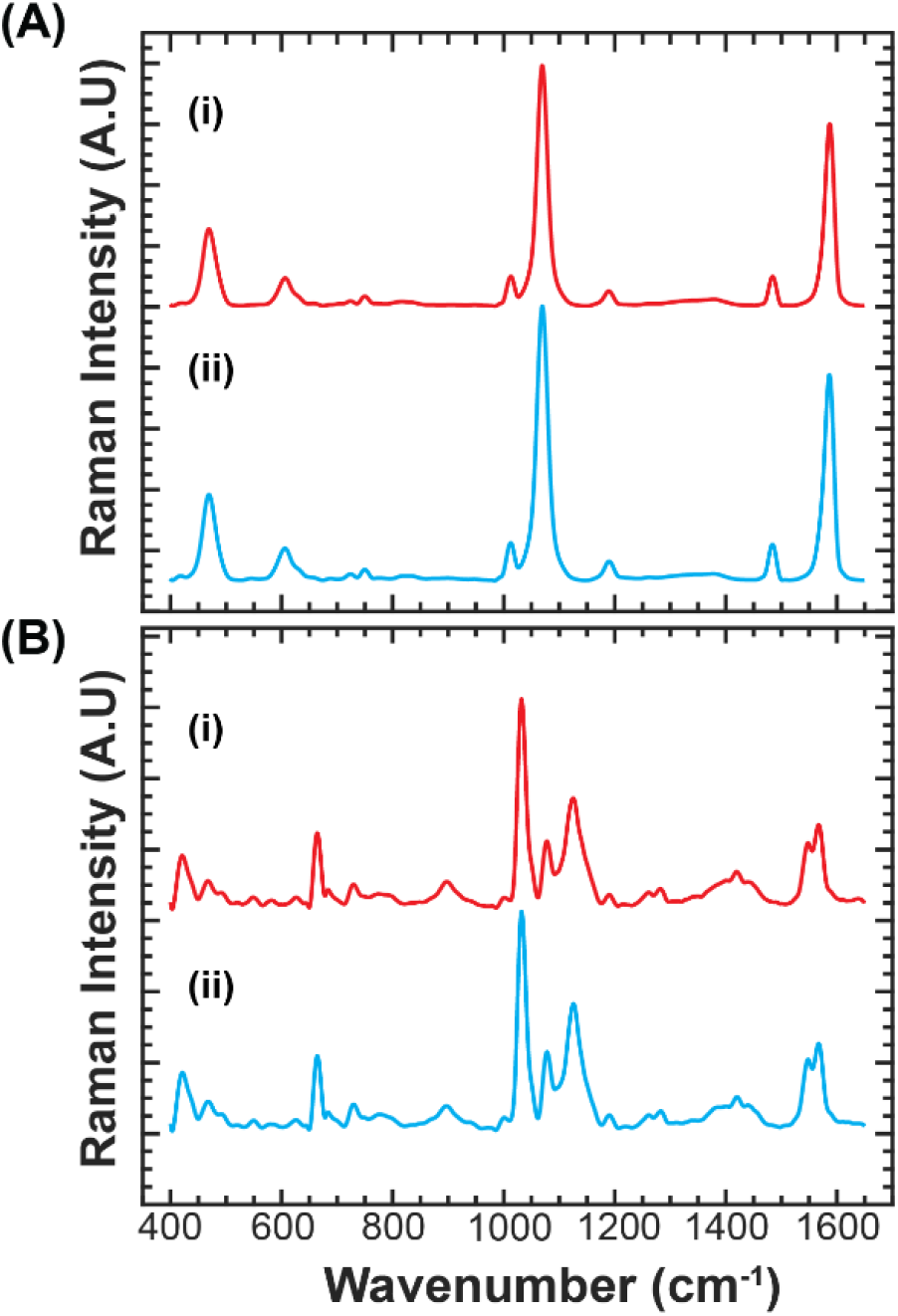
4MBA (A) and 1TBA (B) linkage identification case average SERS spectra. The spectra are lactose (i) and glucose galactose equal molar concentration mixture (ii).

### Enhancing classification accuracy using the collective SERS spectra

4MBA functionalized nanopapers show a good performance in distinguishing the seven selected monosaccharides, maltose/isomaltose, and lactose. 1TBA functionalized nanopapers failed to distinguish lactose but showed an acceptable accuracy for classifying the other glycans. When the number of the classification groups increases, the computation complexity of machine learning classifier increases dramatically^42^. To evaluate the limitation of the machine learning classifier, we analyzed all of the spectra previously collected from 11 sample groups, including 7 monosaccharides, maltose, isomaltose, lactose, and the mixture of galactose and glucose. (**Fig. S11**). Although the classification performance is still acceptable after increasing the number of classification groups, the decreases in the average classification accuracies for both 4MBA and 1TBA were 0.8% and 8.8%, respectively.

As discussed above, glycan binding to different boronic acids could induce changes in their vibration modes, offering different structural information. We hypothesized that the collective SERS spectra from different boronic acid receptors could enrich the structural information, eventually improving the classification accuracy. This multi-receptor concept for SERS detection has been implemented to improve the accuracy of beverage classification^43, 44^. To evaluate this strategy, we integrated the SERS spectra collected from 4MBA and 1TBA functionalized nanopapers and analyzed the collective spectra using the same machine learning algorithm. The confusion matrix of the collective spectra classification is shown in **Fig. S12**. The major variations of the collective spectra could be observed by PCA plot (**Fig. S13~S15**). In PC1 (explained 38.88% variation), the major spectral variations were contributed by 4MBA. The information from the 1TBA nanopaper was dominated in PC2 (explained by 31.29% variation). Compared with the average classification accuracies obtained using the spectra from an individual boronic acid receptor, 99.2% for 4MBA and 91.2% for 1TBA, we observed an improvement in average classification accuracy, 99.6%, from the collective spectra.

### Quantification of sialic acids

We demonstrated that the combination of boronic acid receptors, SERS, and machine learning algorithms could classify the selected glycan samples. In addition to qualitative analysis, we inquired whether the sensing platform could offer quantitative analysis. To evaluate the potential of glycan quantification, we measured the mole fractions of sialic acids.

Sialic acids play essential roles in cellular communication and the immune system. The amount of sialic acids in milk is critical for infant health^45^. For example, goat milk has a relatively higher sialic acid content than bovine milk products^45, 46^; as such, goat milk is considered as a better alternative to human breast milk. A quick and easy way to monitor the sialic acid concentration could assist in the quality control of dairy products.

Since lactose is highly abundant in milk, we mixed sialic acids with lactose to mimic the milk glycosylation environment. The mixtures of sialic acids and lactose were spotted on 4MBA and 1TBA functionalized nanopapers, and the SERS spectra were collected as previously. The spectra were processed by DAPC, and the canonical variables were used to build a machine learning regression model. We chose Gaussian process regression (GPR), a nonparametric regression model, because GPR has been successfully used in material science, chemistry, bacteria identifications, and milk processing quality control^47–51^. We established the GPR models using the SERS spectra from a single boronic acid (4MBA or 1TBA) and the collective spectra. The mole fractions of sialic acids spiked in the background lactose range from 0 mol.% to 2mol.%. The quantification model performance was evaluated by 5-fold cross-validation.

4MBA shows a better quantification performance than 1TBA. (**Fig. S16 & S17**) The 4MBA model showed good prediction (R^2^ of 0.9978) and a low NMSE (0.00214). 1TBA model still exhibited an acceptable quantification performance (R^2^ of 0.9937 and NMSE of 0.00629). The performance of quantification could be further improved using the collective spectra. (**Fig. S18**) The model based on the collective spectra could reach the best R^2^ of 0.9981 and the lowest NMSE of 0.00195.

The data demonstrated that this sensing platform is capable of quantifying sialic acids within background glycans. The performance of this sensor is satisfactory for the quality control of goat milk products. The prior study measured the sialic acid concentration of goat milk during different lactation periods^46^. The concentrations of sialic acids in goat milk for lactation periods I to IV are 46.5, 63.1, 50.0, 12.2 mg/100 ml goat milk, and the mole fractions of sialic acid against lactose ranged from 0.3 mol.% to 1.5 mol.%^46^. Based on the regression result (**Fig. S18**), the model is sufficient to quantify the mole fractions of sialic acid in goat milk.

## Conclusion

We have designed a machine learning-driven SERS glycan sensor that enables the classification of the selected glycan with over 99% accuracy. The boronic acid receptors (4MBA and 1TBA) effectively capture glycan molecules through the selective cis-diol chemical interactions. The DFT simulations show that glycan-boronic acid reactions alter the molecular vibrations and charge distributions, inducing distinct spectral variations. Even stereoisomers (glucose, mannose, and galactose) could trigger sufficient spectral changes that can be detected by the machine learning algorithm. Furthermore, the glycan bindings can influence the vibrations of the aromatic ring structures on the boronic acid receptors; therefore, each boronic acid receptor could offer a unique glycan binding spectrum. By integrating the spectra obtained from 4MBA and 1TBA receptors, the structural information of the glycan was enriched, leading to improved classification accuracy.

Recognition of isomeric structures is one of the major challenges in glycan analysis^2^. This SERS glycan sensor not only distinguishes the stereoisomers (glucose, mannose, and galactose) but also the structural isomers with different glycosidic linkages. The sensor could recognize α-1,4 and α-1,6 glycosidic linkages in maltose and isomaltose, as well as the β-1,4 glycosidic linkage between glucose and galactose in lactose molecules. In addition to qualitative classification, this sensor can quantify sialic acids within the lactose background. The sensor can detect the mole fractions of sialic acids in lactose using the machine learning regression model. The quantification accuracy was further improved by using the collective spectra obtained from 4MBA and 1TBA receptors. The detection accuracy is sufficient to differentiate the amounts of sialic acids in goat milk during different lactation periods^46^.

In summary, the combination of SERS, boronic receptors, and machine learning-driven chemometrics offers a rapid analytical approach for glycan detection. The collective spectra from multiple types of boronic acid receptors can enrich the structural information of glycans, leading to improved classification and quantification accuracies. This study focused on the analysis of small oligosaccharides for proof-of-concept. For more complex glycan structures, the detection accuracy may be improved by using an array of boronic acid receptors.

## Supporting Information

Movie 1~3: the simulated vibration modes and corresponding peaks (MP4)

Peak assignments for 4MBA and 1TBA, Glycan and boronic acid structure, Scheme for machine learning, SERS spectra for monosaccharides, electrostatic potential distribution from DFT simulation, PCA contribution plots and confusion matrix for classification cases, regression model performance plots (PDF)

## Acknowledgements

The authors gratefully acknowledge the support from National Science Foundation (CHE-1904784 and CBET-2114203) and the Human Resource Development Program for Industrial Innovation funded by the Korea Institute for Advancement of Technology (KIAT). Computational resources from the Texas A&M University High Performance Research Computing are gratefully acknowledged.

## References

(1) Gagneux, P.; Hennet, T.; Varki, A. Biological Functions of Glycans. Essentials of Glycobiology [Internet]. 4th edition 2022.

(2) Cummings, R. D.; Pierce, J. M. The challenge and promise of glycomics. Chem Biol 2014, 21 (1), 1–15. DOI: 10.1016/j.chembiol.2013.12.010

(3) Marino, K.; Bones, J.; Kattla, J. J.; Rudd, P. M. A systematic approach to protein glycosylation analysis: a path through the maze. Nat Chem Biol 2010, 6 (10), 713–723. DOI: 10.1038/nchembio.437

(4) Mimura, Y.; Katoh, T.; Saldova, R.; O’Flaherty, R.; Izumi, T.; Mimura-Kimura, Y.; Utsunomiya, T.; Mizukami, Y.; Yamamoto, K.; Matsumoto, T.; et al. Glycosylation engineering of therapeutic IgG antibodies: challenges for the safety, functionality and efficacy. Protein Cell 2018, 9 (1), 47–62. DOI: 10.1007/s13238-017-0433-3

(5) Zhang, X.; Dai, Z.; Si, S.; Zhang, X.; Wu, W.; Deng, H.; Wang, F.; Xiao, X.; Jiang, C. Ultrasensitive SERS Substrate Integrated with Uniform Subnanometer Scale “Hot Spots” Created by a Graphene Spacer for the Detection of Mercury Ions. Small 2017, 13 (9), 1603347. DOI: 10.1002/smll.201603347.

(6) Weatherston, J. D.; Seguban, R. K. O.; Hunt, D.; Wu, H. J. Low-Cost and Simple Fabrication of Nanoplasmonic Paper for Coupled Chromatography Separation and Surface Enhanced Raman Detection. ACS Sens 2018, 3 (4), 852–857. DOI: 10.1021/acssensors.8b00098

(7) Weatherston, J. D.; Yuan, S.; Mashuga, C. V.; Wu, H. J. Multi-functional SERS substrate: collection, separation, and identification of airborne chemical powders on a single device. Sens Actuators B Chem 2019, 297, 126765. DOI: 10.1016/j.snb.2019.126765

(8) Weatherston, J. D.; Worstell, N. C.; Wu, H. J. Quantitative surface-enhanced Raman spectroscopy for kinetic analysis of aldol condensation using Ag-Au core-shell nanocubes. Analyst 2016, 141 (21), 6051–6060, 10.1039/C6AN01098A. DOI: 10.1039/c6an01098a

(9) Hu, Q.; Sellers, C.; Kwon, J. S.-I.; Wu, H.-J. Integration of surface-enhanced Raman spectroscopy (SERS) and machine learning tools for coffee beverage classification. Digital Chemical Engineering 2022, 3, 100020. DOI: 10.1016/j.dche.2022.100020.

(10) Wang, X.; Xia, N.; Liu, L. Boronic Acid-based approach for separation and immobilization of glycoproteins and its application in sensing. Int J Mol Sci 2013, 14 (10), 20890–20912. DOI: 10.3390/ijms141020890

(11) Deng, R.; Yue, J.; Qu, H.; Liang, L.; Sun, D.; Zhang, J.; Liang, C.; Xu, W.; Xu, S. Glucose-bridged silver nanoparticle assemblies for highly sensitive molecular recognition of sialic acid on cancer cells via surface-enhanced raman scattering spectroscopy. Talanta 2018, 179, 200–206. DOI: 10.1016/j.talanta.2017.11.006

(12) Liang, L.; Qu, H.; Zhang, B.; Zhang, J.; Deng, R.; Shen, Y.; Xu, S.; Liang, C.; Xu, W. Tracing sialoglycans on cell membrane via surface-enhanced Raman scattering spectroscopy with a phenylboronic acid-based nanosensor in molecular recognition. Biosens Bioelectron 2017, 94, 148–154. DOI: 10.1016/j.bios.2017.02.043

(13) Li, S.; Zhou, Q.; Chu, W.; Zhao, W.; Zheng, J. Surface-enhanced Raman scattering behaviour of 4-mercaptophenyl boronic acid on assembled silver nanoparticles. Phys Chem Chem Phys 2015, 17 (27), 17638–17645. DOI: 10.1039/c5cp02409a

(14) Tommasone, S.; Allabush, F.; Tagger, Y. K.; Norman, J.; Kopf, M.; Tucker, J. H. R.; Mendes, P. M. The challenges of glycan recognition with natural and artificial receptors. Chem Soc Rev 2019, 48 (22), 5488–5505. DOI: 10.1039/c8cs00768c

(15) Miyazaki, T.; Khan, T.; Tachihara, Y.; Itoh, M.; Miyazawa, T.; Suganami, T.; Miyahara, Y.; Cabral, H.; Matsumoto, A. Boronic Acid Ligands Can Target Multiple Subpopulations of Pancreatic Cancer Stem Cells via pH-Dependent Glycan-Terminal Sialic Acid Recognition. ACS Appl Bio Mater 2021, 4 (9), 6647–6651. DOI: 10.1021/acsabm.1c00383

(16) Hu, C.; Peng, F.; Mi, F.; Wang, Y.; Geng, P.; Pang, L.; Ma, Y.; Li, G.; Li, Y.; Guan, M. SERS-based boronate affinity biosensor with biomimetic specificity and versatility: Surface-imprinted magnetic polymers as recognition elements to detect glycoproteins. Anal Chim Acta 2022, 1191, 339289. DOI: 10.1016/j.aca.2021.339289

(17) Usta, D. D.; Salimi, K.; Pinar, A.; Coban, I.; Tekinay, T.; Tuncel, A. A Boronate Affinity-Assisted SERS Tag Equipped with a Sandwich System for Detection of Glycated Hemoglobin in the Hemolysate of Human Erythrocytes. ACS Appl Mater Interfaces 2016, 8 (19), 11934–11944. DOI: 10.1021/acsami.6b00138

(18) Lin, L.; Tian, X.; Hong, S.; Dai, P.; You, Q.; Wang, R.; Feng, L.; Xie, C.; Tian, Z. Q.; Chen, X. A bioorthogonal Raman reporter strategy for SERS detection of glycans on live cells. Angew Chem Int Ed Engl 2013, 52 (28), 7266–7271. DOI: 10.1002/anie.201301387

(19) Tabatabaei, M.; Wallace, G. Q.; Caetano, F. A.; Gillies, E. R.; Ferguson, S. S. G.; Lagugne-Labarthet, F. Controlled positioning of analytes and cells on a plasmonic platform for glycan sensing using surface enhanced Raman spectroscopy. Chem Sci 2016, 7 (1), 575–582. DOI: 10.1039/c5sc03332b

(20) Sun, X.; Zhai, W.; Fossey, J. S.; James, T. D. Boronic acids for fluorescence imaging of carbohydrates. Chem Commun (Camb) 2016, 52 (17), 3456–3469. DOI: 10.1039/c5cc08633g

(21) Yuan, S.; Jiao, Z. R.; Quddus, N.; Kwon, J. S. I.; Mashuga, C. V. Developing Quantitative Structure-Property Relationship Models To Predict the Upper Flammability Limit Using Machine Learning. Industrial & Engineering Chemistry Research 2019, 58 (8), 3531–3537. DOI: 10.1021/acs.iecr.8b05938.

(22) Yuan, S.; Zhang, Z.; Sun, Y.; Kwon, J. S.; Mashuga, C. V. Liquid flammability ratings predicted by machine learning considering aerosolization. J Hazard Mater 2020, 386, 121640. DOI: 10.1016/j.jhazmat.2019.121640

(23) Lussier, F.; Thibault, V.; Charron, B.; Wallace, G. Q.; Masson, J. F. Deep learning and artificial intelligence methods for Raman and surface-enhanced Raman scattering. Trac-Trends in Analytical Chemistry 2020, 124, 115796. DOI: 10.1016/j.trac.2019.115796.

(24) Jombart, T.; Devillard, S.; Balloux, F. Discriminant analysis of principal components: a new method for the analysis of genetically structured populations. BMC Genet 2010, 11 (1), 94. DOI: 10.1186/1471-2156-11-94

(25) Berrar, D. Cross-Validation. In Encyclopedia of Bioinformatics and Computational Biology, Ranganathan, S., Gribskov, M., Nakai, K., Schönbach, C. Eds.; Academic Press, 2019; pp 542–545. DOI: 10.1016/b978-0-12-809633-8.20349-x

(26) Trevethan, R. Sensitivity, Specificity, and Predictive Values: Foundations, Pliabilities, and Pitfalls in Research and Practice. Front Public Health 2017, 5 (307), 307, Perspective. DOI: 10.3389/fpubh.2017.00307

(27) Krishnan, R.; Binkley, J. S.; Seeger, R.; Pople, J. A. Self-consistent molecular orbital methods. XX. A basis set for correlated wave functions. J. Chem. Phys. 1980, 72 (1), 650–654. DOI: 10.1063/1.438955.

(28) Grimme, S.; Antony, J.; Ehrlich, S.; Krieg, H. A consistent and accurate ab initio parametrization of density functional dispersion correction (DFT-D) for the 94 elements H-Pu. J. Chem. Phys. 2010, 132 (15), 154104. DOI: 10.1063/1.3382344.

(29) National Institute of Standards and Technology, https://www.nist.gov/.

(30) Revanna, B. N.; Madegowda, M. Dithiane Based Boronic Acid as a Carbohydrate Sensor in an Aqueous Solution at pH 7.5: Theoretical and Experimental Approach. J Fluoresc 2021, 31 (6), 1683–1703. DOI: 10.1007/s10895-021-02791-4

(31) Su, H.; Wang, Y.; Yu, Z.; Liu, Y.; Zhang, X.; Wang, X.; Sui, H.; Sun, C.; Zhao, B. Surface-enhanced Raman spectroscopy study on the structure changes of 4-Mercaptophenylboronic Acid under different pH conditions. Spectrochim Acta A Mol Biomol Spectrosc 2017, 185, 336–342. DOI: 10.1016/j.saa.2017.05.068

(32) Jamroz, M. H. Vibrational energy distribution analysis (VEDA): scopes and limitations. Spectrochim Acta A Mol Biomol Spectrosc 2013, 114, 220–230. DOI: 10.1016/j.saa.2013.05.096

(33) Sharma, B.; Frontiera, R. R.; Henry, A. I.; Ringe, E.; Van Duyne, R. P. SERS: Materials, applications, and the future. Materials Today 2012, 15 (1-2), 16–25. DOI: 10.1016/S1369-7021(12)70017-2.

(34) Gu, X.; Trujillo, M. J.; Olson, J. E.; Camden, J. P. SERS Sensors: Recent Developments and a Generalized Classification Scheme Based on the Signal Origin. Annu Rev Anal Chem (Palo Alto Calif) 2018, 11 (1), 147–169. DOI: 10.1146/annurev-anchem-061417-125724

(35) Parlak, C.; Ramasami, P.; Tursun, M.; Rhyman, L.; Kaya, M. F.; Atar, N.; Alver, O.; Senyel, M. 4-Mercaptophenylboronic acid: conformation, FT-IR, Raman, OH stretching and theoretical studies. Spectrochim Acta A Mol Biomol Spectrosc 2015, 144, 131–138. DOI: 10.1016/j.saa.2015.02.040

(36) Axthelm, J.; Askes, S. H. C.; Elstner, M.; G, U. R.; Gorls, H.; Bellstedt, P.; Schiller, A. Fluorinated Boronic Acid-Appended Pyridinium Salts and (19)F NMR Spectroscopy for Diol Sensing. J Am Chem Soc 2017, 139 (33), 11413–11420. DOI: 10.1021/jacs.7b01167

(37) Meiland, M.; Heinze, T.; Guenther, W.; Liebert, T. Seven-membered ring boronates at trans-diol moieties of carbohydrates. Tetrahedron Letters 2009, 50 (4), 469–472. DOI: 10.1016/j.tetlet.2008.11.043

(38) Benner, K.; Klufers, P.; Labisch, O. Borate esters of the methyl D-glucopyranosides. Carbohydr Res 2007, 342 (18), 2801–2806. DOI: 10.1016/j.carres.2007.08.022

(39) Griffin, M. E.; Hsieh-Wilson, L. C. Tools for mammalian glycoscience research. Cell 2022, 185 (15), 2657–2677. DOI: 10.1016/j.cell.2022.06.016

(40) Pham, X. H.; Seong, B.; Hahm, E.; Huynh, K. H.; Kim, Y. H.; Kim, J.; Lee, S. H.; Jun, B. H. Glucose Detection of 4-Mercaptophenylboronic Acid-Immobilized Gold-Silver Core-Shell Assembled Silica Nanostructure by Surface Enhanced Raman Scattering. Nanomaterials (Basel) 2021, 11 (4), 948. DOI: 10.3390/nano11040948

(41) Ajit Varki, R. D. C., Jeffrey D. Esko, Pamela Stanley, Gerald W. Hart, Markus Aebi, Debra Mohnen, Taroh Kinoshita, Nicolle H. Packer, James H. Prestegard, Ronald L. Schnaar, and Peter H. Seeberger. Essentials of Glycobiology. 4 ed.; Cold Spring Harbor Laboratory Press: Cold Spring Harbor (NY), 2022.

(42) Ray, S. An Analysis of Computational Complexity and Accuracy of Two Supervised Machine Learning Algorithms—K-Nearest Neighbor and Support Vector Machine. In Data Management, Analytics and Innovation, Singapore, 2021; Sharma, N., Chakrabarti, A., Balas, V. E., Martinovic, J., Eds.; Springer Singapore: pp 335–347.

(43) Li, F.; Wang, X. Q.; Zhou, S. N.; Wang, D. M.; Gong, Z. J.; Fan, M. K. Multidimensional Surface-Enhanced Raman Scattering (SERS) Strategy for Tea Differentiation. Acs Food Science & Technology 2022, 2 (7), 1096–1102. DOI: 10.1021/acsfoodscitech.2c00091

(44) Leong, Y. X.; Lee, Y. H.; Koh, C. S. L.; Phan-Quang, G. C.; Han, X.; Phang, I. Y.; Ling, X. Y. Surface-Enhanced Raman Scattering (SERS) Taster: A Machine-Learning-Driven Multireceptor Platform for Multiplex Profiling of Wine Flavors. Nano Lett 2021, 21 (6), 2642–2649. DOI: 10.1021/acs.nanolett.1c00416

(45) van Leeuwen, S. S.; Te Poele, E. M.; Chatziioannou, A. C.; Benjamins, E.; Haandrikman, A.; Dijkhuizen, L. Goat Milk Oligosaccharides: Their Diversity, Quantity, and Functional Properties in Comparison to Human Milk Oligosaccharides. J Agric Food Chem 2020, 68 (47), 13469–13485. DOI: 10.1021/acs.jafc.0c03766

(46) de Sousa, Y. R. F.; da Silva Vasconcelos, M. A.; Costa, R. G.; de Azevedo Filho, C. A.; de Paiva, E. P.; Queiroga, R. d. C. R. d. E. Sialic acid content of goat milk during lactation. Livestock Science 2015, 177, 175–180. DOI: 10.1016/j.livsci.2015.04.005.

(47) Schulz, E.; Speekenbrink, M.; Krause, A. A tutorial on Gaussian process regression: Modelling, exploring, and exploiting functions. Journal of Mathematical Psychology 2018, 85, 1–16. DOI: 10.1016/j.jmp.2018.03.001.

(48) Deringer, V. L.; Bartok, A. P.; Bernstein, N.; Wilkins, D. M.; Ceriotti, M.; Csanyi, G. Gaussian Process Regression for Materials and Molecules. Chem Rev 2021, 121 (16), 10073–10141. DOI: 10.1021/acs.chemrev.1c00022

(49) Kemmler, M.; Rodner, E.; Rosch, P.; Popp, J.; Denzler, J. Automatic identification of novel bacteria using Raman spectroscopy and Gaussian processes. Anal Chim Acta 2013, 794, 29–37. DOI: 10.1016/j.aca.2013.07.051

(50) Kemmler, M.; Denzler, J.; Rösch, P.; Popp, J. Classification of Microorganisms via Raman Spectroscopy Using Gaussian Processes. In Pattern Recognition, Berlin, Heidelberg, 2010//, 2010; Goesele, M., Roth, S., Kuijper, A., Schiele, B., Schindler, K., Eds.; Springer Berlin Heidelberg: pp 81–90.

(51) Vasafi, P. S.; Hinrichs, J.; Hitzmann, B. Establishing a novel procedure to detect deviations from standard milk processing by using online Raman spectroscopy. Food Control 2022, 131, 108442. DOI: 10.1016/j.foodcont.2021.108442.

