## Supplementary information for "Advancing Glycan Analysis: A New Platform Integrating SERS, Boronic Acids, and Machine Learning Algorithms"

Table S1: Peak assignment for 4MBA………………………......………………………………..... Page 2

Table S2: Peak assignment for 1TBA……………………………………………..…………...…..… Page 3

Figure S1: Glycan and boronic acid structure……….…………………………………………... Page 4

Figure S2: Scheme for machine learning……………………………..……………..………………Page 5

Figure S3: SERS spectra for monosaccharides……………………………………………………...Page 6

Figure S4: Molecular electrostatic potential distribution for 1TBA and 1TBA-glycan simulation........…...Page 7

Figure S5: PCA contribution plot for monosaccharide classification….……………………….……...Page 8

Figure S6: Confusion matrix for monosaccharide classification…………………………………..…Page 10

Figure S7: PCA contribution plot for disaccharide classification………..........................………..…...Page 11

Figure S8: Confusion matrix for disaccharide classification…………………………………......…...Page 13

Figure S9: PCA contribution plot for β-1,4 linkages classification………..………………………………Page 14

Figure S10: Confusion matrix for β-1,4 linkages classification……………………………..……..…...Page 16

Figure S11: Confusion matrix for combined cases…………………………………………………...Page 17

Figure S12: Confusion matrix for collective spectra……………..…………………………………..Page 18

Figure S13: PC1 contribution plot for collective spectra……………………….…………...…………..Page 19

Figure S14: PC2 contribution plot for collective spectra………………………………...……………..Page 19

Figure S15: PC3 contribution plot for collective spectra……………...……………………………...…...Page 20

Figure S16: Regression model performance for 4MBA spectra……………………………..…….Page 21

Figure S17: Regression model performance for 1TBA spectra…………………………..…….……….Page 21

Figure S18: Regression model performance for collective spectra……………..…………………………Page 22

**Table S1. Peak assignment for 4MBA**

| SERS Frequency ($cm^{-1}$) | Scaled Calculated Frequency ($cm^{-1}$) | Vibrational Assignment | Reference |
|---|---|---|---|
| 470-476 | 460 | CCC out of plane bending | [1, 2] |
| 607-611 | 590 | CCC in plane bending | [1, 2] |
| 641-644 | 634 | CS stretching | [1, 2] |
| 720 | 720 | CCC in-plane bending | [1-3] |
| 750 | 743 | CH stretching | [1, 2] |
| 894 | 882 | CSH in plane bending | [2, 4] |
| 1008-1010 | 1008 | CC stretching, OH stretching | [1, 2, 4] |
| 1027-1035 | 1040 | CC stretching, OH stretching | [4] |
| 1057-1060 | 1061 | CC stretching, OH stretching | [1, 2, 4] |
| 1074-1078 | 1079 | CCC in plane bending, CS stretching | [1, 2] |
| 1187-1190 | 1193 | CH stretching, BOH in plane bending | [1, 2, 4] |
| 1221 | 1226 | CH stretching | [1] |
| 1472-1474 | 1452 | CC stretching, BC stretching | [1, 2, 4] |
| 1488 | 1518 | CC stretching, BC stretching | [1, 2, 4] |
| 1570 | 1560 | CC stretching, CH bending | [1, 2, 4] |
| 1589 | 1610 | CC stretching, CH bending | [1, 2, 4] |

**Table S2. Peak assignment for 1TBA**

| SERS Frequency ($cm^{-1}$) | Scaled Calculated Frequency ($cm^{-1}$) | Vibrational Assignment (PED≥5%) | Reference |
|---|---|---|---|
| 418 | 406 | SC stretching, SCC in-plane bending | [5] |
| 434 | 428 | CCCC torsion, SCCC out-of-plane bending | [5] |
| 460 | 459 | SCC torsion, CSCC torsion | [5] |
| 477 | 486 | CCCC torsion, CSCC torsion, HOBC torsion, CSCC out-of-plane bending, BCCC out-of-plane bending, OCOB out-of-plane bending | [5] |
| 493 | 518 | SCC in-plane bending | [5] |
| 627 | 594 | CCCC torsion, CCSC torsion, HOBC torsion, CSCC out-of-plane bending, BCCC out-of-plane bending, CCCS out-of-plane bending | [5] |
| 664 | 663 | CCCC torsion (64), CCC in-plane bending (14) | VEDA |
| 679 | 673 | CCC in-plane bending | [5] |
| 713 | 719 | SCCC out-of-plane bending (29), CCCC torsion (27) | VEDA |
| 729 | 727 | CCC in plane bending (31), HOBC torsion (11) | VEDA |
| 755 | 740 | CCCC torsion (44), CCC in-plane bending (28), SC stretching (22) | VEDA |
| 775 | 774 | HCCC torsion | [5] |
| 921 | 925 | HCCC torsion (78) | VEDA |
| 999 | 992 | OB stretching, HOB in-plane bending | [5] |
| 1018 | 1022 | CCS in-plane bending, CCC in-plane bending | [5] |
| 1031 | 1038 | CC stretching | [5] |
| 1046 | 1050 | CCC in-plane bending (25) | VEDA |
| 1078 | 1087 | CC stretching (64) | VEDA |
| 1096 | 1099 | CC stretching (24) | VEDA |
| 1118 | 1114 | CC stretching (43), HCC in-plane bending (16) | VEDA |
| 1134 | 1132 | HCC in-plane bending (21), CC stretching (13) | VEDA |
| 1190 | 1191 | HCC in-plane bending (72), CC stretching (26) | VEDA |
| 1223 | 1210 | CC stretching (33) | VEDA |
| 1283 | 1259 | CC stretching | [5] |
| 1421 | 1430 | HCC in-plane bending (56), CC stretching (21) | VEDA |
| 1442 | 1447 | HCC in-plane bending (57), CC stretching (14) | VEDA |
| 1456 | 1455 | HCC in-plane bending (52), CCC in-plane bending (17) | VEDA |
| 1548 | 1544 | CC stretching | [5] |
| 1567 | 1571 | CC stretching (30) | VEDA |
| 1583-1585 | 1580 | CC stretching (29), HCC in-plane bending (22) | VEDA |

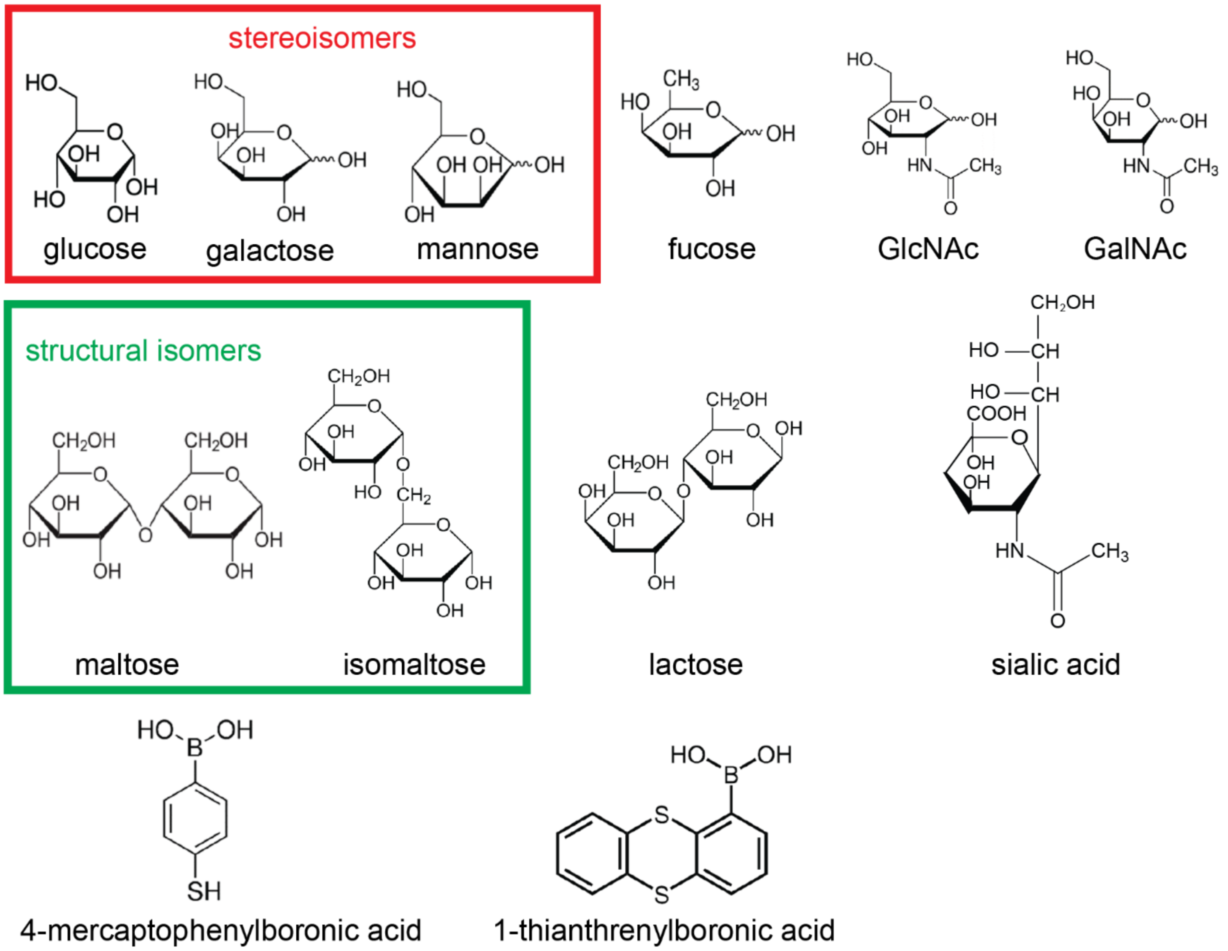


**Figure S1.** Boronic acid and glycan structures

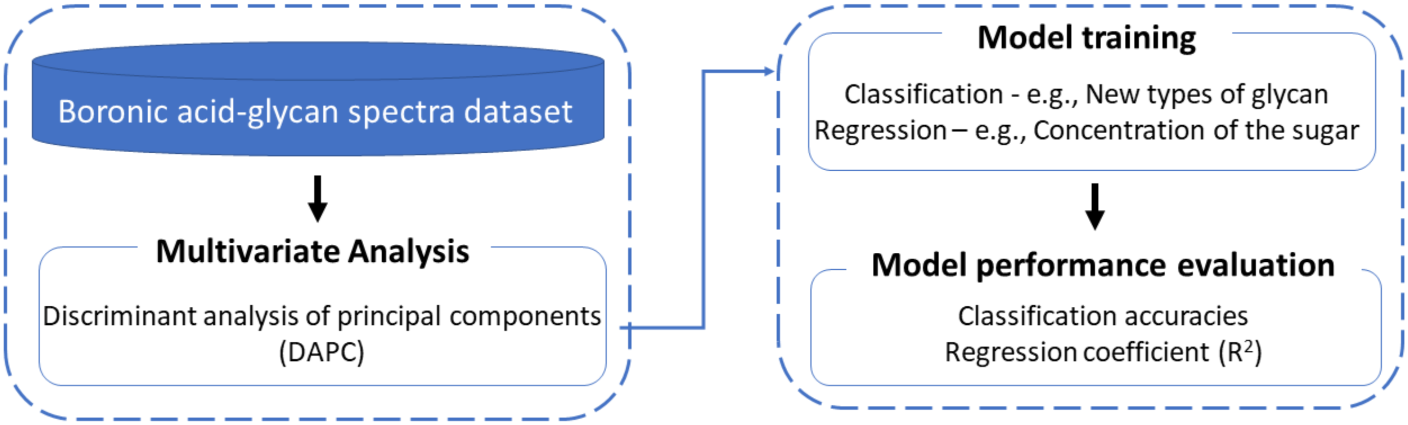


**Figure S2.** Scheme for machine learning

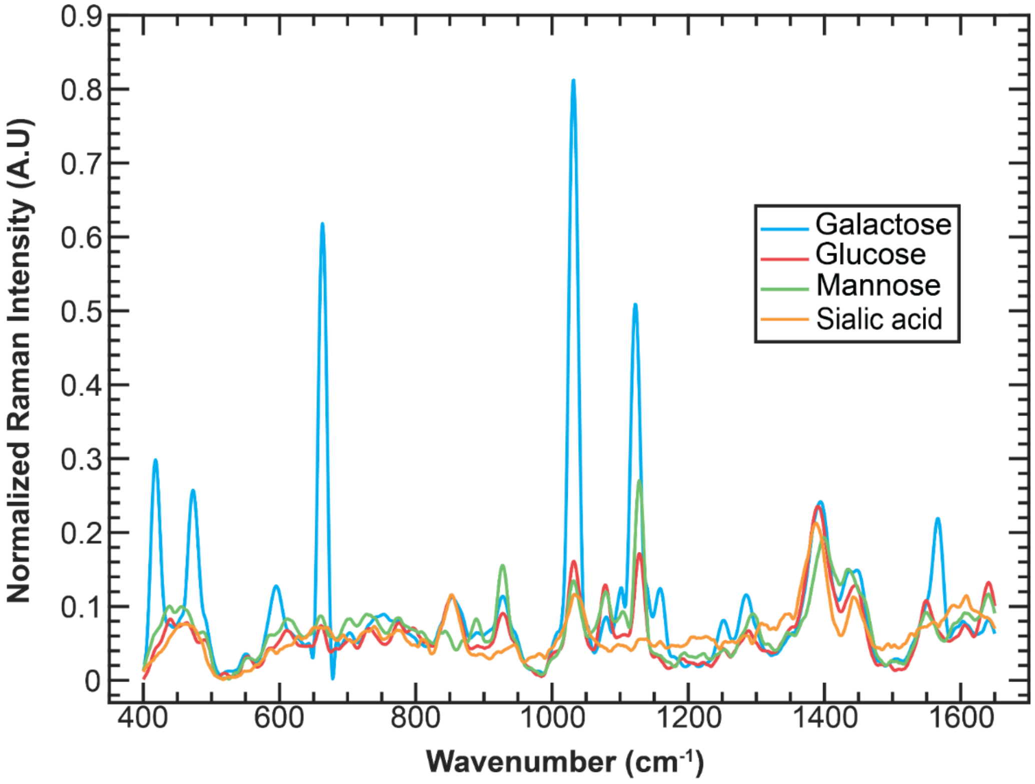


**Figure S3.** SERS spectra of monosaccharides on bare nanopaper (no boronic acid functionalization). The peaks overlap and cause difficulties in interpretation.

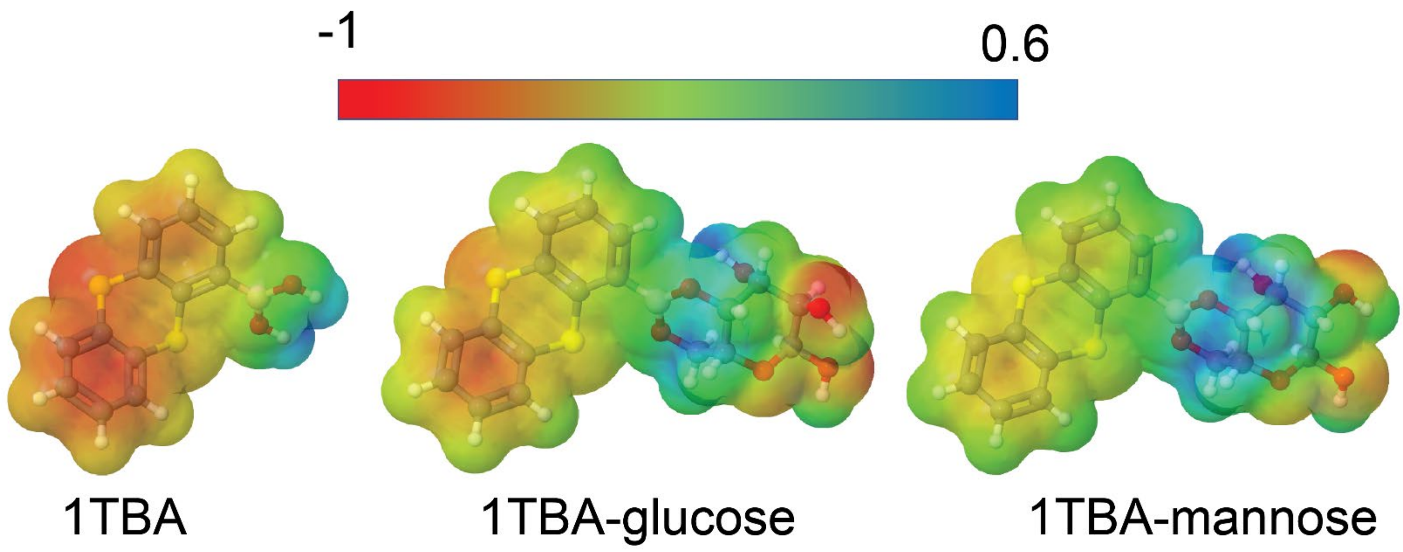


**Figure S4.** Molecular electrostatic potential distribution for 1TBA, 1TBA-glucose, and 1TBA-mannose. The color shows the calculated Mulliken charges carried by the atoms. Change distribution changes on the aromatic ring structure of the 1TBA when 1TBA bind with the glucose and mannose.

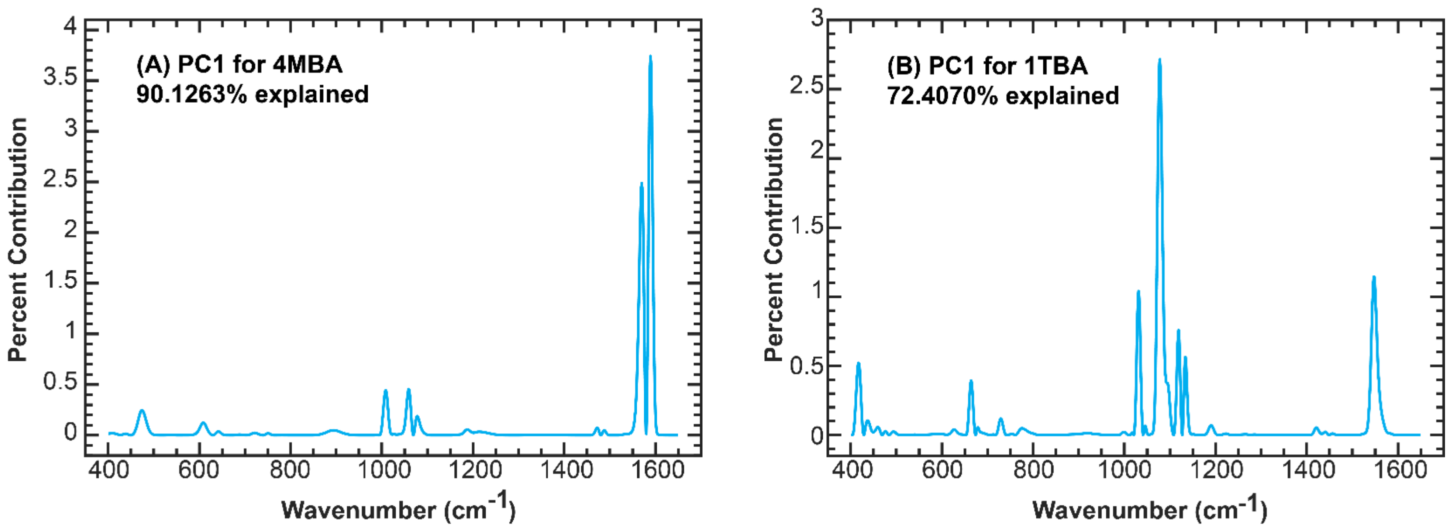


**Figure S5.** PCA contribution plot for PC1 in monosaccharide case using 4MBA (A) (90.1263% explained) and 1TBA (B) (72.4070% explained). The tables below show the peaks appears in the contribution plot and corresponding to **Table S1** and **S2**.

**Peak assignment for 4MBA monosaccharide PC1 contribution plot**

| Frequency ($cm^{-1}$) | Vibrational Assignment |
|---|---|
| 472 | CCC out-of-plane bending |
| 607 | CCC in-plane bending |
| 642 | CS stretching |
| 721 | CCC in-plane bending |
| 751 | CH stretching |
| 895 | CSH in-plane bending |
| 1008 | CC stretching, OH stretching |
| 1059 | CC stretching, OH stretching |
| 1078 | CCC in-plane bending, CS stretching |
| 1187 | CH stretching, BOH in-plane bending |
| 1472 | CC stretching, BC stretching |
| 1488 | CC stretching, BC stretching |
| 1570 | CC stretching, CH bending |
| 1589 | CC stretching, CH bending |

**Peak assignment for 1TBA monosaccharide PC1 contribution plot**

| Frequency ($cm^{-1}$) | Vibrational Assignment |
|---|---|
| 418 | SC stretching, SCC in-plane bending |
| 437 | CCCC torsion, SCCC out-of-plane bending |
| 460 | SCC torsion, CSCC torsion |
| 476 | CCCC torsion, CSCC torsion, HOBC torsion, CSCC out-of-plane bending, BCCC out-of-plane bending, OCOB out-of-plane bending |
| 494 | SCC in-plane bending |
| 626 | CCCC torsion, CCSC torsion, HOBC torsion, CSCC out-of-plane bending, BCCC out-of-plane bending, CCCS out-of-plane bending |
| 664 | CCCC torsion, CCC in-plane bending |

| | |
|---|---|
| 678 | CCC in-plane bending |
| 713 | SCCC out-of-plane bending, CCCC torsion |
| 728 | CCC in-plane bending, HOBC torsion |
| 755 | CCC in-plane bending, CCCC torsion, SC stretching |
| 775 | HCCC torsion |
| 921 | HCCC torsion |
| 999 | OB stretching, HOB in-plane bending |
| 1018 | CCS in-plane bending, CCC in-plane bending |
| 1031 | CC stretching |
| 1046 | CCC in-plane bending |
| 1078 | CC stretching |
| 1092 | CC stretching |
| 1119 | CC stretching, HCC in-plane bending |
| 1134 | HCC in-plane bending, CC stretching |
| 1190 | HCC in-plane bending, CC stretching |
| 1223 | CC stretching |
| 1420 | HCC in-plane bending, CC stretching |
| 1442 | HCC in-plane bending, CC stretching |
| 1457 | HCC in-plane bending, CCC in-plane bending |
| 1548 | CC stretching |
| 1583 | CC stretching, HCC in-plane bending |

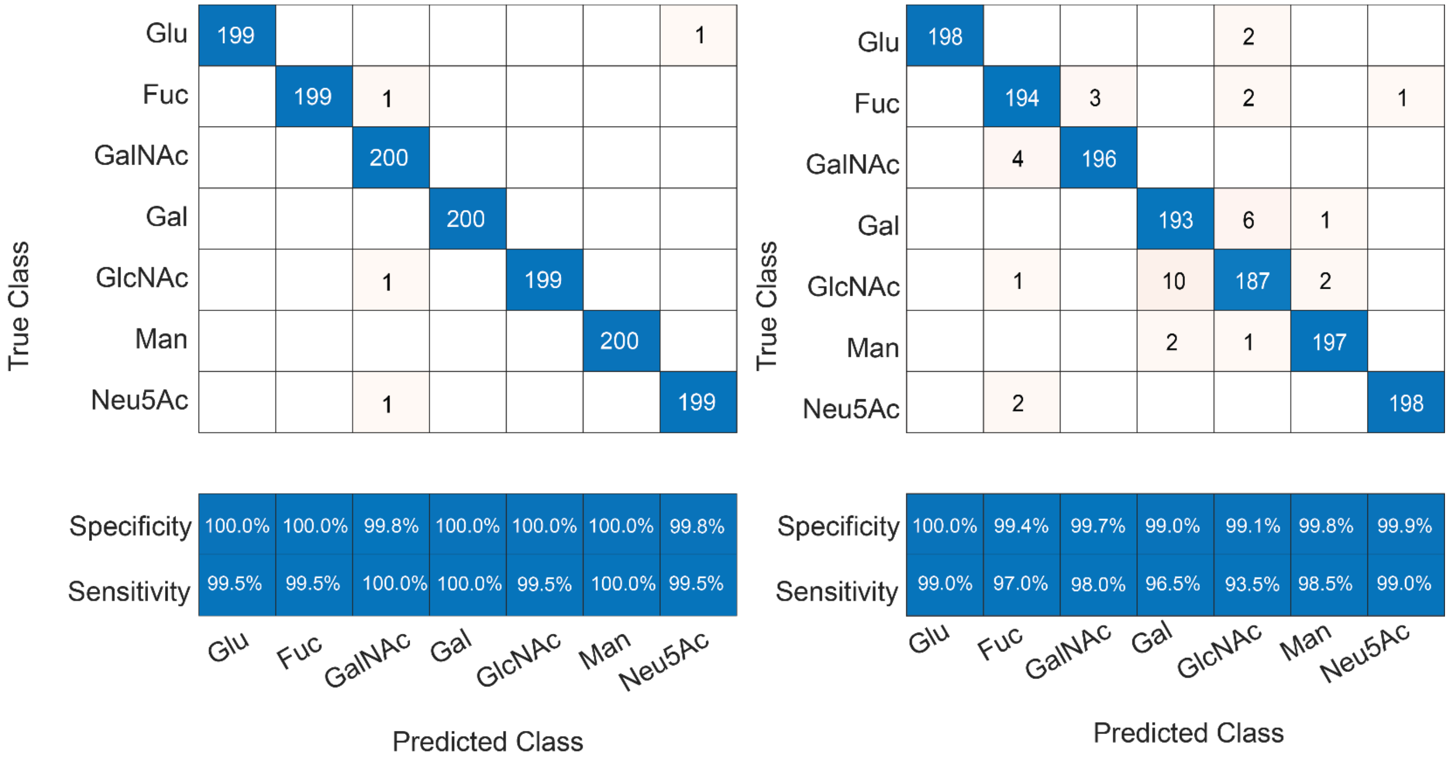


**Figure S6.** Confusion matrix for monosaccharide classification for 4MBA (A) (99.7% accuracy) and 1TBA (B) (97.4% accuracy).

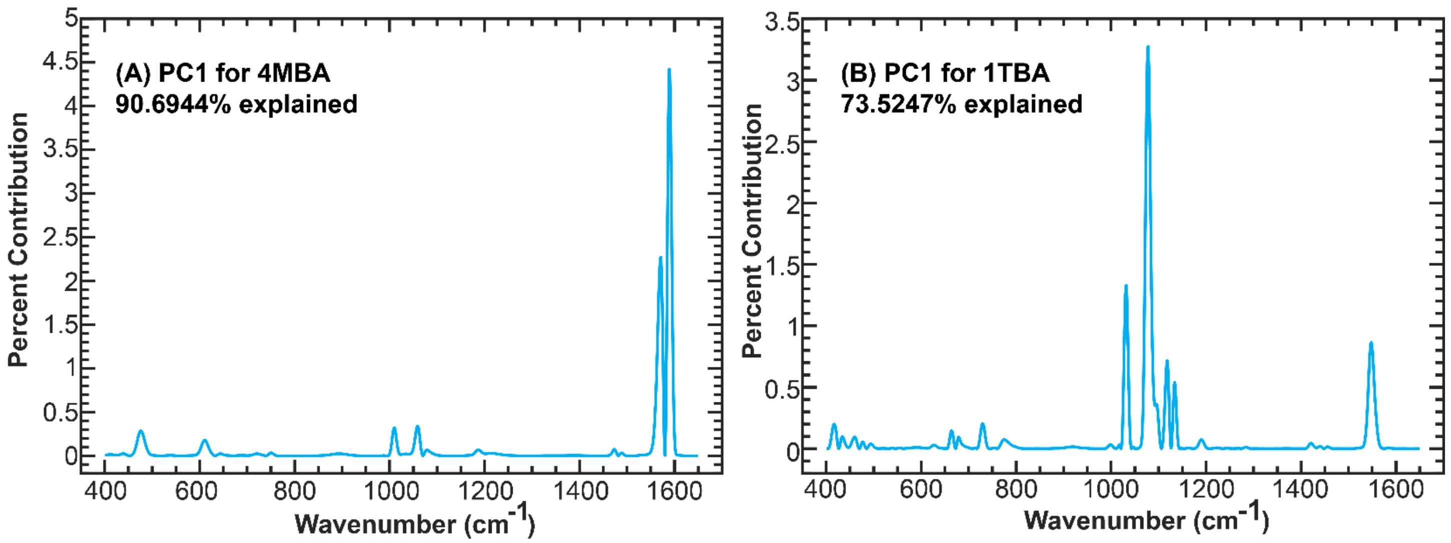


**Figure S7** PCA contribution plot for PC1 in disaccharide case using 4MBA (A) (90.6944% explained) and 1TBA (B) (73.5247% explained). The tables below show the peaks appear in the contribution plot and corresponding to **Table S1** and **S2**.

**Peak assignment for 4MBA disaccharide PC1 contribution plot**

| Frequency ($cm^{-1}$) | Vibrational Assignment |
|---|---|
| 476 | CCC out-of-plane bending |
| 611 | CCC in-plane bending |
| 644 | CS stretching |
| 720 | CCC in-plane bending |
| 750 | CH stretching |
| 895 | CSH in-plane bending |
| 1010 | CC stretching, OH stretching |
| 1035 | CC stretching, OH stretching |
| 1059 | CC stretching, OH stretching |
| 1078 | CCC in-plane bending, CS stretching |
| 1187 | CH stretching, BOH in-plane bending |
| 1474 | CC stretching, BC stretching |
| 1489 | CC stretching, BC stretching |
| 1571 | CC stretching, CH bending |
| 1589 | CC stretching, CH bending |

**Peak assignment for 1TBA disaccharide PC1 contribution plot**

| Frequency ($cm^{-1}$) | Vibrational Assignment |
|---|---|
| 417 | SC stretching, SCC in-plane bending |
| 434 | CCCC torsion, SCCC out-of-plane bending |
| 460 | SCC torsion, CSCC torsion |
| 477 | CCCC torsion, CCSC torsion, CCSC out-of-plane bending, BCCC out-of-plane bending |
| 493 | SCC in-plane bending |
| 627 | CCCC torsion, CCSC torsion, HOBC torsion, CSCC out-of-plane bending, BCCC out-of-plane bending, CCCS out-of-plane bending |

| | |
|---|---|
| 664 | CCCC torsions, CCC in-plane bending |
| 679 | CCC in-plane bending |
| 713 | SCCC out-of-plane bending, CCCC torsion |
| 729 | CCC in-plane bending, HOBC torsion |
| 754 | CCC in-plane bending, CCCC torsion, SC stretching |
| 775 | HCCC torsion |
| 920 | HCCC torsion |
| 998 | OB stretching, HOB in-plane bending |
| 1017 | CCS in-plane bending, CCC in-plane bending |
| 1031 | CC stretching |
| 1046 | CCC in-plane bending |
| 1078 | CC stretching |
| 1096 | CC stretching |
| 1118 | CC stretching, HCC in-plane bending |
| 1134 | HCC in-plane bending, CC stretching |
| 1189 | HCC in-plane bending, CC stretching |
| 1283 | CC stretching |
| 1421 | HCC in-plane bending, CC stretching |
| 1440 | HCC in-plane bending, CC stretching |
| 1456 | HCC in-plane bending, CCC in-plane bending |
| 1548 | CC stretching |
| 1585 | CC stretching, HCC in-plane bending |

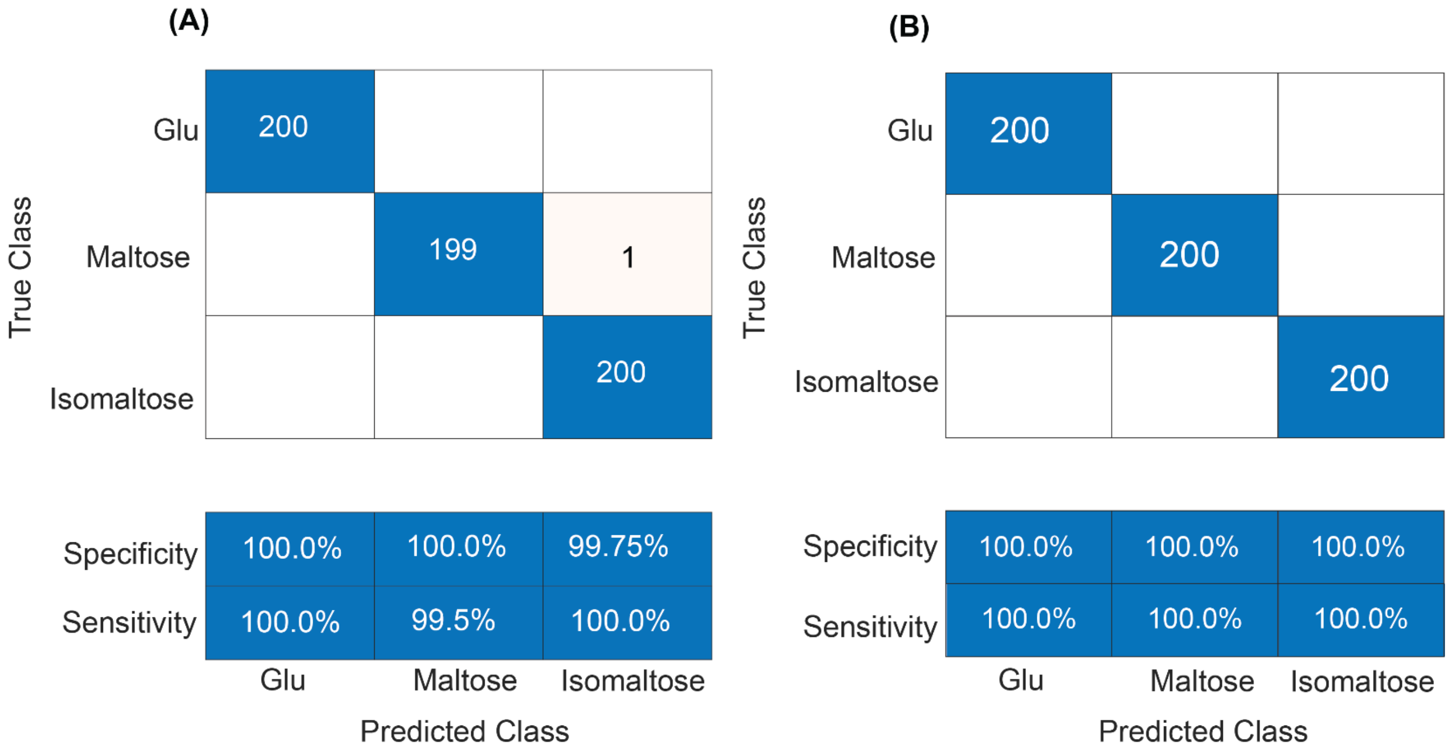


**Figure S8.** Confusion matrix for disaccharide case using 4MBA (A) (99.8% accuracy) and 1TBA (B) (100% accuracy)

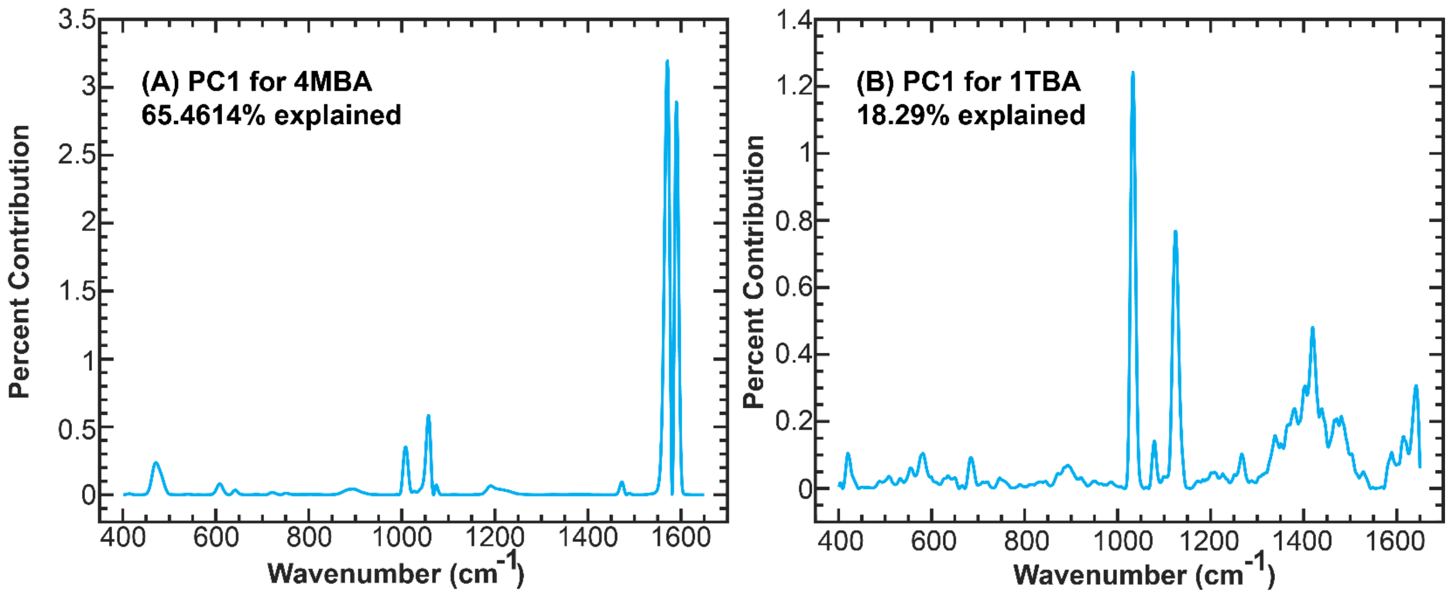


**Figure S9.** PCA contribution plot for identifying β-1,4 glycosidic linkage in lactose using 4MBA (A) (65.4614% explained) and 1TBA (B) (18.29% explained). The tables below show the peaks appear in the contribution plot and corresponding to **Table S1** and **S2**.

**Peak assignment for 4MBA links PC1 contribution plot**

| Frequency ($cm^{-1}$) | Vibrational Assignment |
|---|---|
| 470 | CCC out-of-plane bending |
| 607 | CCC in-plane bending |
| 641 | CS stretching |
| 720 | CCC in-plane bending |
| 751 | CH stretching |
| 894 | CSH in-plane bending |
| 1008 | CC stretching, OH stretching |
| 1027 | CC stretching, OH stretching |
| 1057 | CC stretching, OH stretching |
| 1074 | CCC in-plane bending, CS stretching |
| 1190 | CH stretching, BOH in-plane bending |
| 1221 | CH stretching |
| 1473 | CC stretching, BC stretching |
| 1489 | CC stretching, BC stretching |
| 1571 | CC stretching, CH bending |
| 1591 | CC stretching, CH bending |

**Peak assignment for 1TBA links PC1 contribution plot**

| Frequency ($cm^{-1}$) | Vibrational Assignment |
|---|---|
| 417 | SC stretching, SCC in-plane bending |
| 434 | CCCC torsion, SCCC out-of-plane bending |
| 460 | SCC torsion, CSCC torsion |
| 477 | CCCC torsion, CCSC torsion, HOBC torsion, CCSC out-of-plane bending, BCCC out-of-plane bending, OCOB out-of-plane bending |
| 493 | SCC in-plane bending |

| | |
|---|---|
| 627 | CCCC torsion, CCSC torsion, HOBC torsion, CSCC out-of-plane bending, BCCC out-of-plane bending, CCCS out-of-plane bending |
| 664 | CCCC torsion, CCC in-plane bending |
| 679 | CCC in-plane bending |
| 713 | SCCC out-of-plane bending, CCCC torsion |
| 729 | CCC in-plane bending, HOBC torsion |
| 754 | CCC in-plane bending, CCCC torsion, SC stretching |
| 775 | HCCC torsion |
| 920 | HCCC torsion |
| 998 | OB stretching, HOB in-plane bending |
| 1017 | CCS in-plane bending, CCC in-plane bending |
| 1031 | CC stretching |
| 1046 | CCC in-plane bending |
| 1078 | CC stretching |
| 1096 | CC stretching |
| 1118 | CC stretching, HCC in-plane bending |
| 1134 | HCC in-plane bending, CC stretching |
| 1189 | HCC in-plane bending, CC stretching |
| 1283 | CC stretching |
| 1421 | HCC in-plane bending, CC stretching |
| 1440 | HCC in-plane bending, CC stretching |
| 1456 | HCC in-plane bending, CCC in-plane bending |
| 1548 | CC stretching |
| 1585 | CC stretching, HCC in-plane bending |

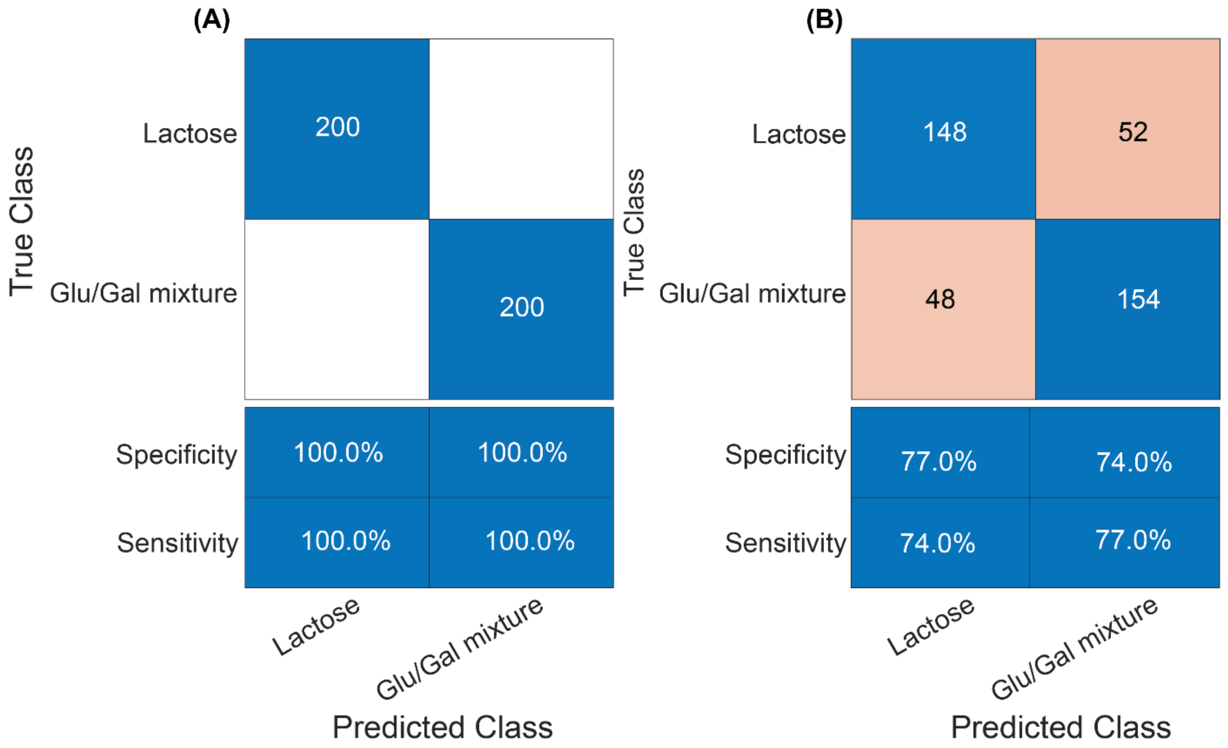


**Figure S10.** Confusion matrix for identifying β-1,4 glycosidic linkage in lactose using 4MBA (A) (100% accuracy) and 1TBA (B) (75.5% accuracy).

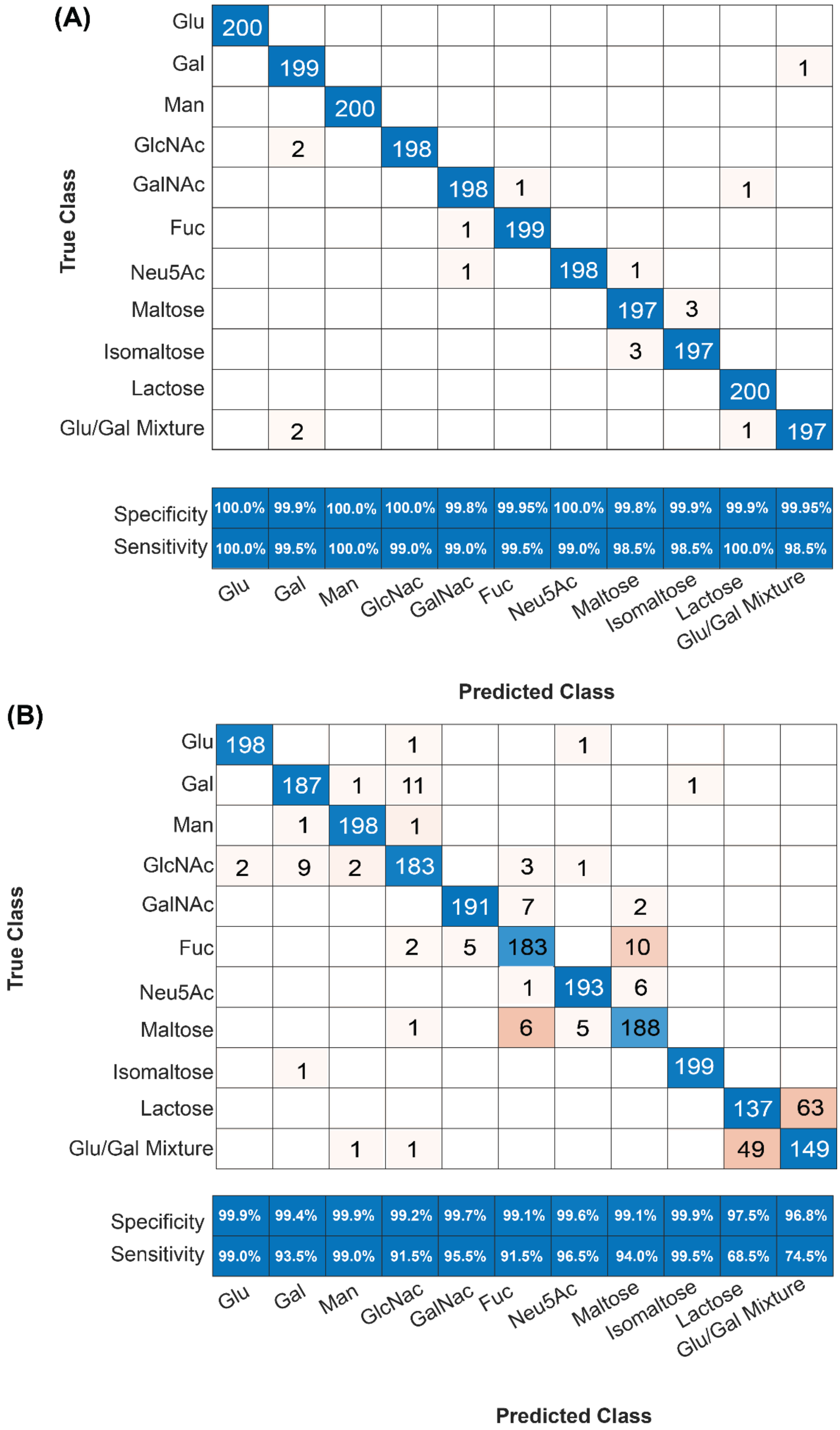


**Figure S11.** Confusion matrix for all studied cases using 4MBA (A) (99.2% accuracy) and 1TBA (B) (91.2% accuracy).

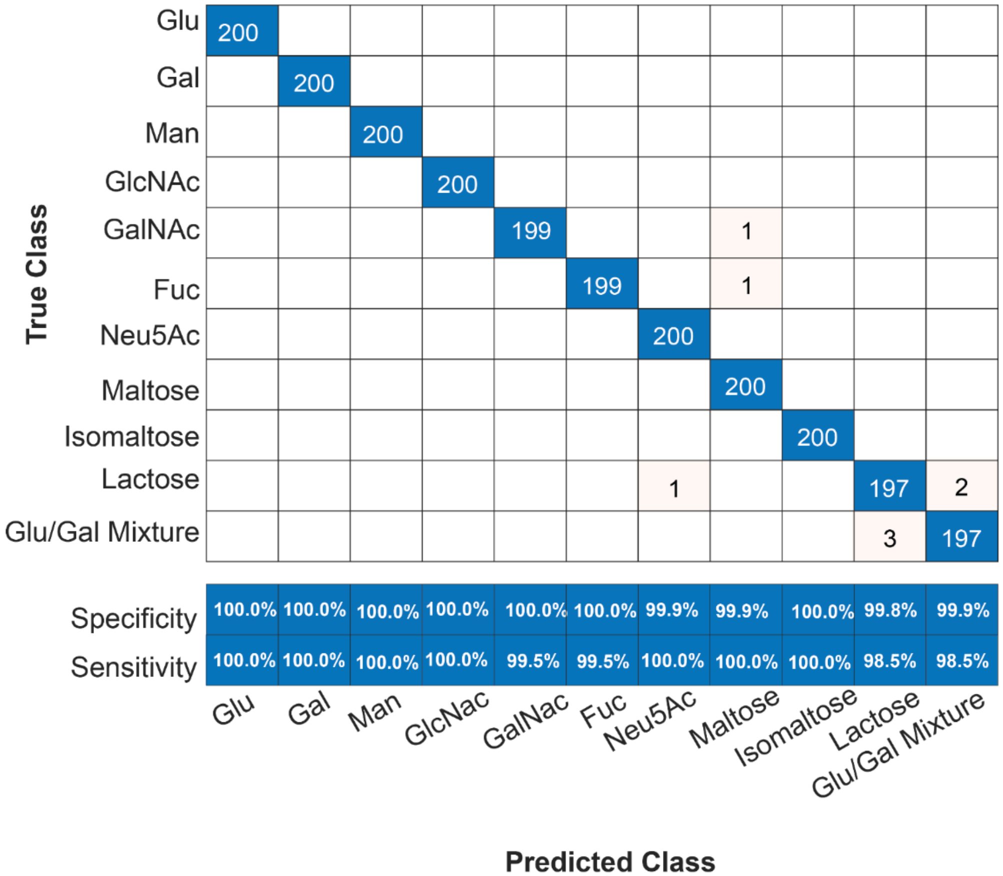


**Figure S12.** Confusion matrix for collective spectra model (99.6% accuracy).

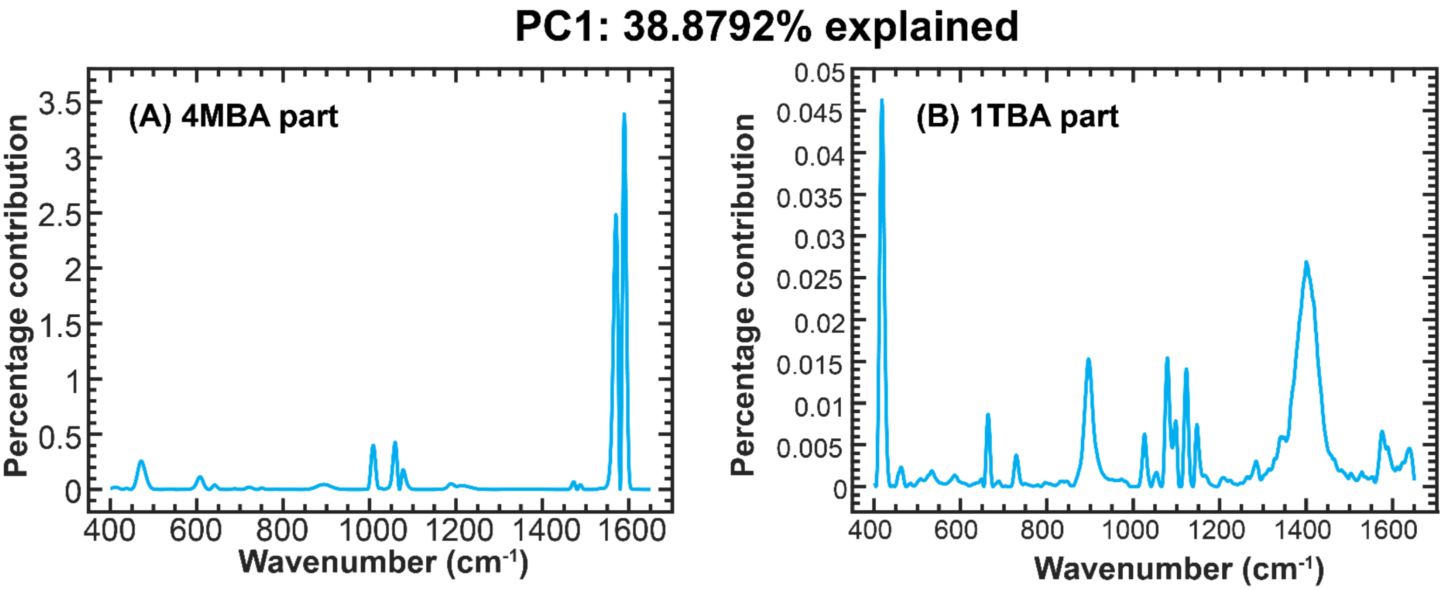


**Figure S13** The percentage contribution from each wavenumber to PC1 (38.8792% explained). The major contribution peaks are from the 4MBA for PC1.

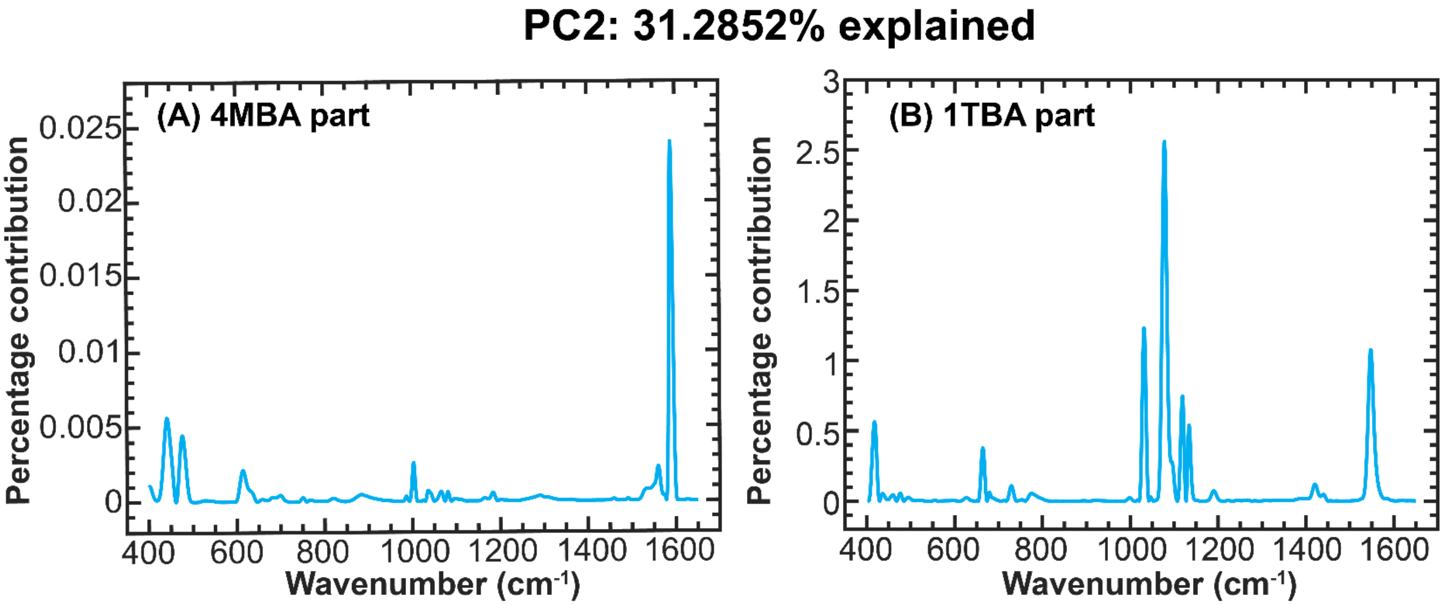


**Figure S14.** The percentage contribution from each wavenumber to PC2 (31.2852% explained). The major contribution peaks are from the 1TBA for PC2.

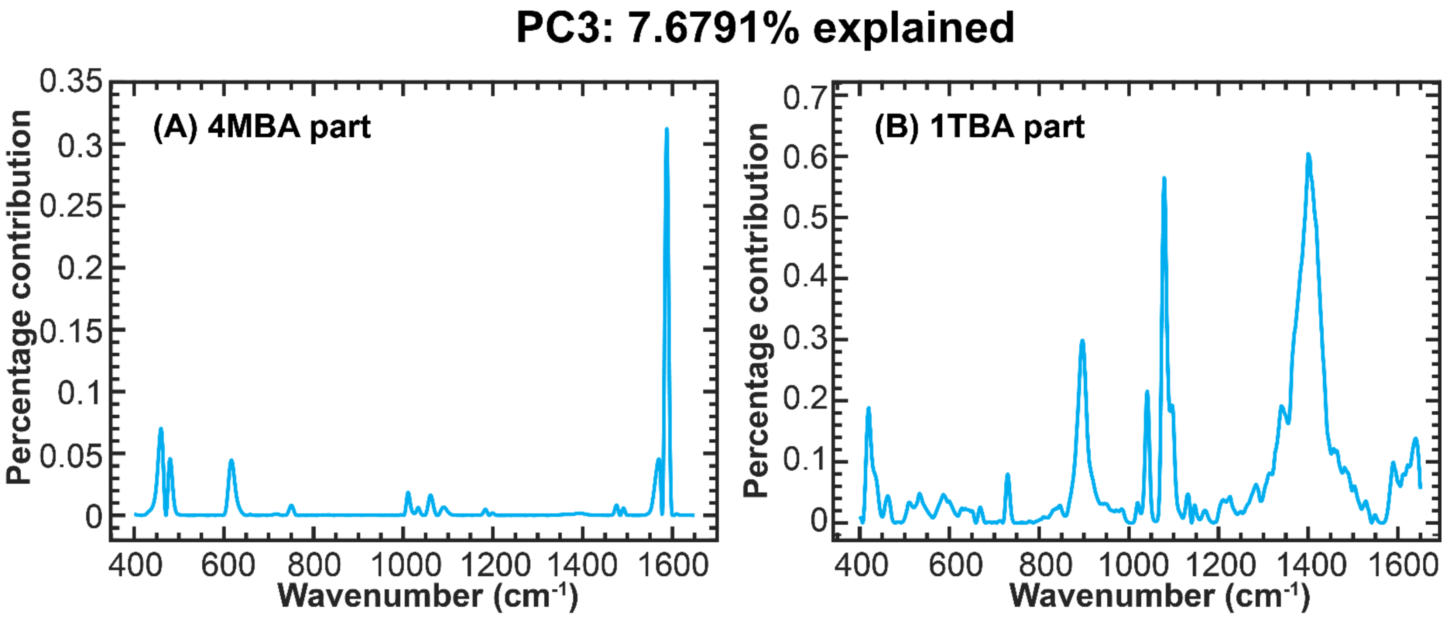


**Figure S15.** The percentage contribution from each wavenumber to PC3 (7.6791% explained). The major contribution peaks are from the both BAs for PC3.

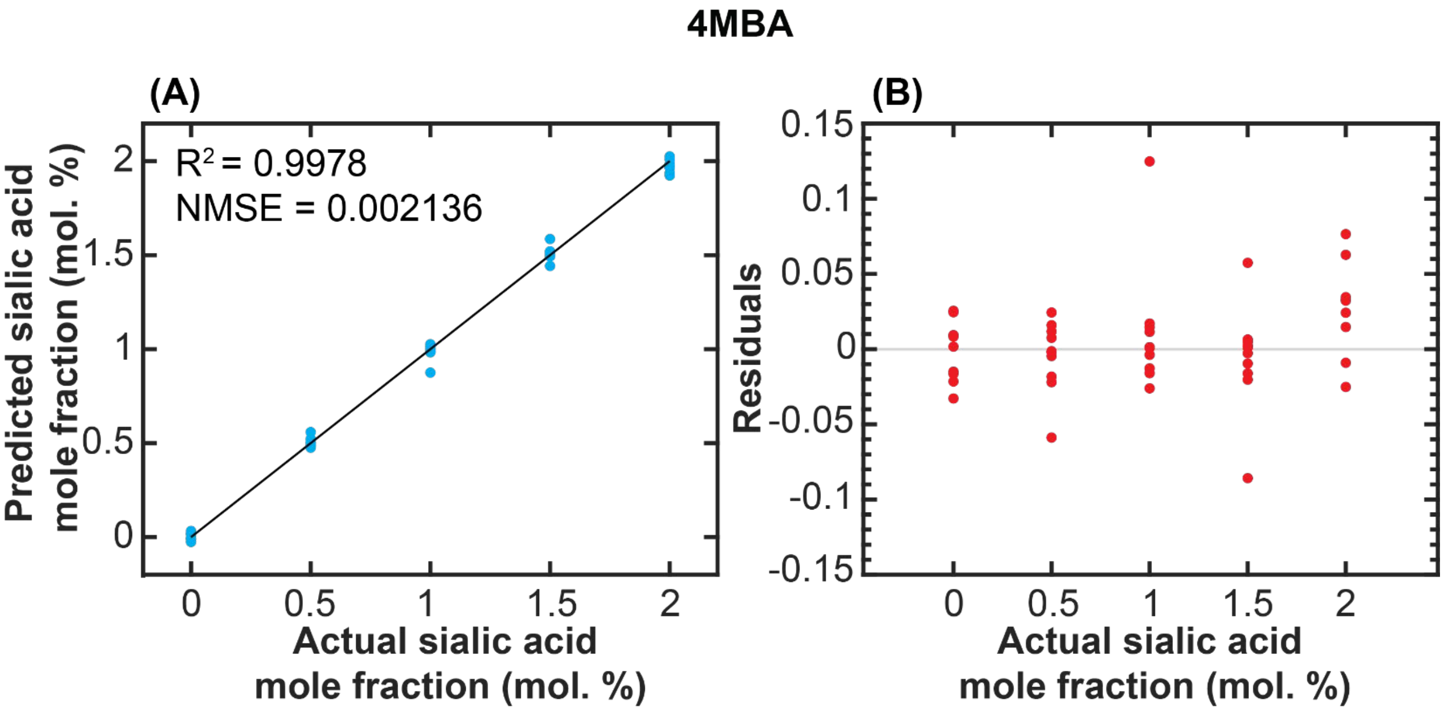


**Figure S16.** Predicted value vs. actual value (A) and residual vs. actual value (B) for 4MBA regression model.

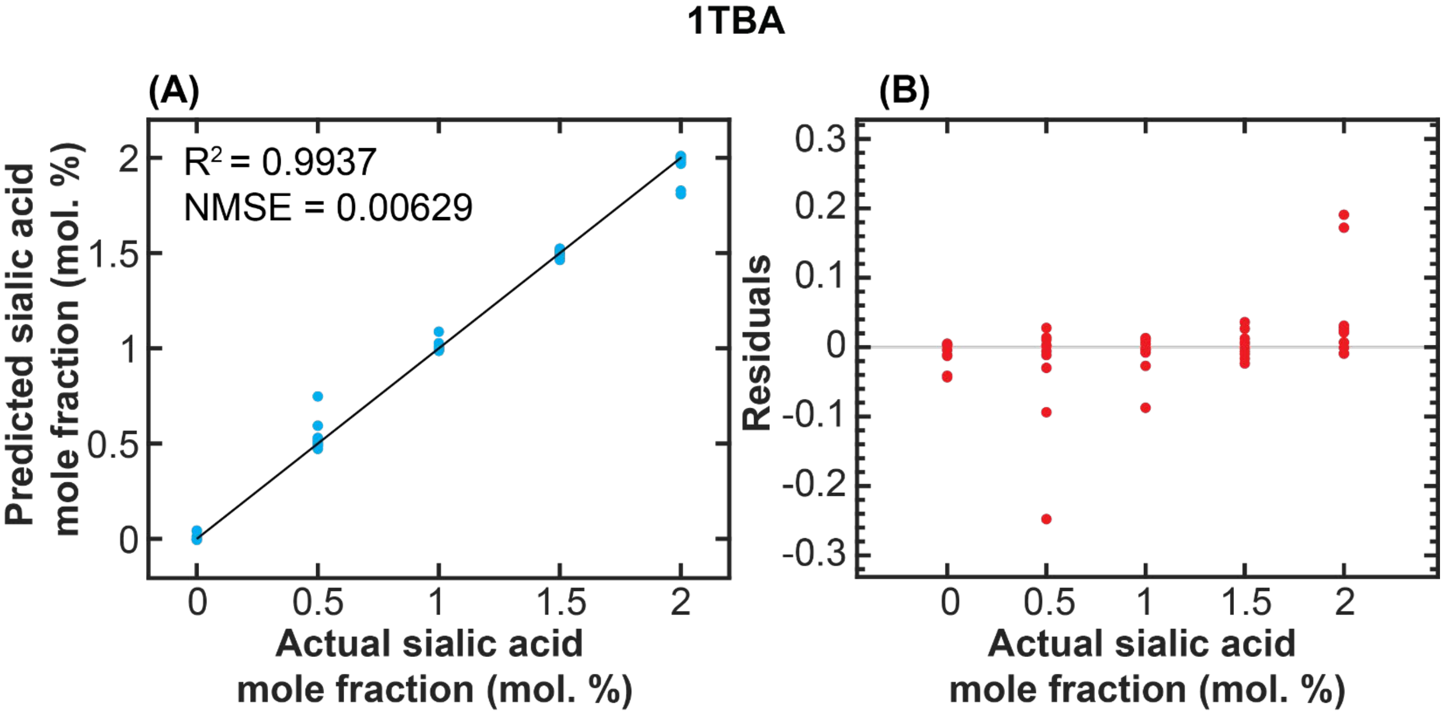


**Figure S17.** Predicted value vs. actual value (A) and residual vs. actual value (B) for 1TBA regression model.

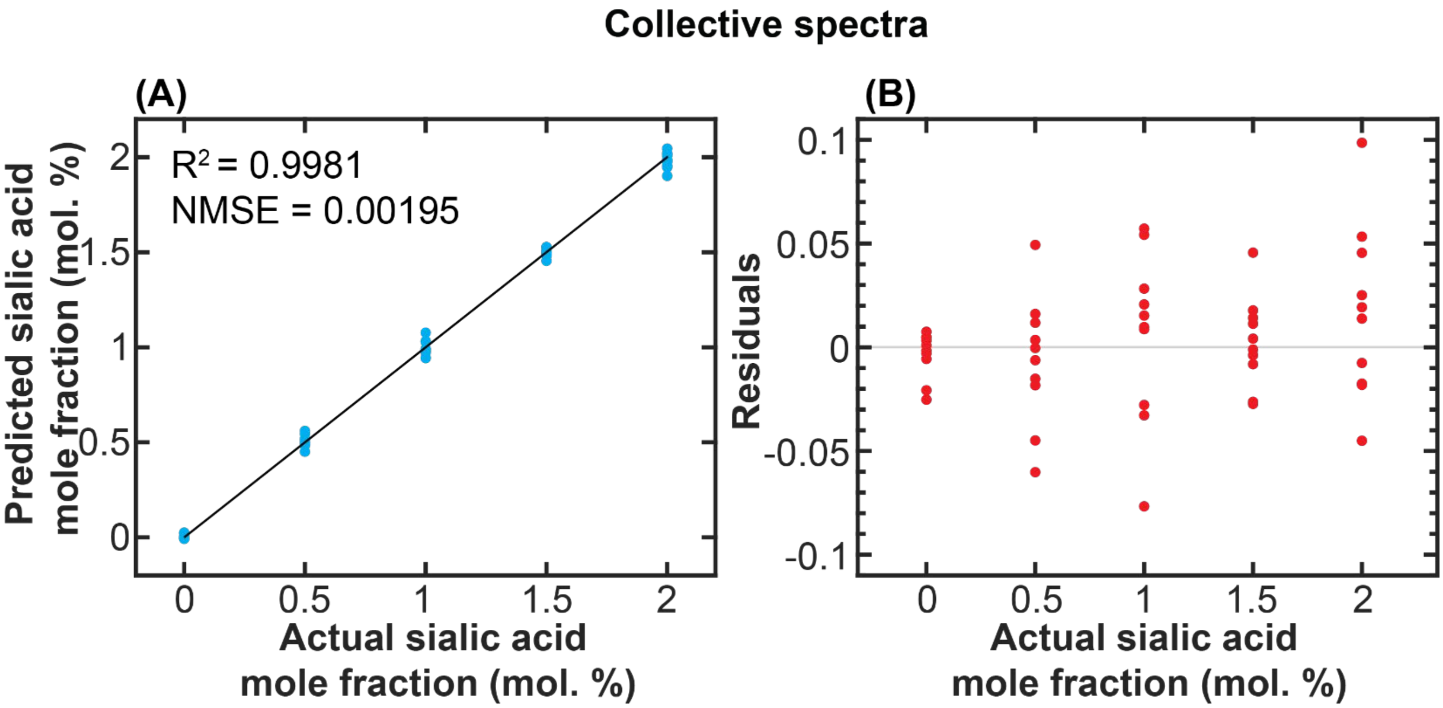


**Figure S18.** Predicted value vs. actual value (A) and residual vs. actual value (B) for collective spectra regression model.

### Reference

(1) Li, S.; Zhou, Q.; Chu, W.; Zhao, W.; Zheng, J. Surface-enhanced Raman scattering behaviour of 4-mercaptophenyl boronic acid on assembled silver nanoparticles. *Phys Chem Chem Phys* **2015**, *17* (27), 17638-17645. DOI: 10.1039/c5cp02409a (acccessed 2021-10-20T19:51:58). From NLM PubMed-not-MEDLINE.

(2) Su, H.; Wang, Y.; Yu, Z.; Liu, Y.; Zhang, X.; Wang, X.; Sui, H.; Sun, C.; Zhao, B. Surface-enhanced Raman spectroscopy study on the structure changes of 4-Mercaptophenylboronic Acid under different pH conditions. *Spectrochim Acta A Mol Biomol Spectrosc* **2017**, *185*, 336-342. DOI: 10.1016/j.saa.2017.05.068 From NLM PubMed-not-MEDLINE.

(3) Liang, L.; Qu, H.; Zhang, B.; Zhang, J.; Deng, R.; Shen, Y.; Xu, S.; Liang, C.; Xu, W. Tracing sialoglycans on cell membrane via surface-enhanced Raman scattering spectroscopy with a phenylboronic acid-based nanosensor in molecular recognition. *Biosens Bioelectron* **2017**, *94*, 148-154. DOI: 10.1016/j.bios.2017.02.043 From NLM Medline.

(4) Parlak, C.; Ramasami, P.; Tursun, M.; Rhyman, L.; Kaya, M. F.; Atar, N.; Alver, O.; Senyel, M. 4-Mercaptophenylboronic acid: conformation, FT-IR, Raman, OH stretching and theoretical studies. *Spectrochim Acta A Mol Biomol Spectrosc* **2015**, *144*, 131-138. DOI: 10.1016/j.saa.2015.02.040 (acccessed 2021-12-16T14:19:57). From NLM Medline.

(5) Revanna, B. N.; Madegowda, M. Dithiane Based Boronic Acid as a Carbohydrate Sensor in an Aqueous Solution at pH 7.5: Theoretical and Experimental Approach. *J Fluoresc* **2021**, *31* (6), 1683-1703. DOI: 10.1007/s10895-021-02791-4 (acccessed 2021-12-16T14:20:18). From NLM PubMed-not-MEDLINE.
